# Ecological patterns of root nodule diversity in cultivated and wild rooibos populations: a community prediction approach

**DOI:** 10.1101/2020.01.15.907972

**Authors:** Josep Ramoneda, Jaco Le Roux, Emmanuel Frossard, Beat Frey, Hannes Andres Gamper

## Abstract

There is interest in understanding the factors behind the biogeography of root-associated bacteria due to the joint effects that plant host, climate, and soil conditions can have on bacterial diversity. For legume crops with remaining wild populations, this is of even more importance, because the effects of cropping on undisturbed root-associated bacterial communities can be addressed. Here, we used a community prediction approach to describe the diversity of the root nodule bacterial communities of rooibos (*Aspalathus linearis*), an endemic legume crop from South Africa. The goal was to reveal whether patterns of root nodule community composition in paired cultivated and wild rooibos populations could be related to geographical distance, plant traits, and plant population type (i.e. cultivated or uncultivated). We identified a core of dominant and widespread *Mesorhizobium* ZOTUs that each defined one of 4 different root nodule community classes. Rooibos cultivation impacted root nodule bacterial diversity at regional and local scales, while the geographical origin of the root nodule communities was the strongest predictor of root nodule community structure. Beyond impacts of cultivation on root nodule bacterial diversity, this study suggests a mixture of dispersal limitation and ecological drift regionally, and selection by different plant populations locally, define the biogeography of rooibos root nodule bacterial communities.

## Introduction

Microbial diversity usually shows strong biogeographic structure, shaped by historical processes and contemporary environmental conditions (Fierer and Jackson, 2006; Hanson *et al.*, 2012). Taxon-area relationships, whereby diversity increases with sample area (Martiny *et al.*, 2006), and distance-decay relationships, whereby communities become more dissimilar with increasing distance, are commonplace for bacteria (Horner-Devine *et al.*, 2004; Zinger *et al.*, 2014; Van Cauwbenberge *et al.*, 2015; Stefan *et al.*, 2018). These ecological patterns result from related, but distinct, ecological processes such as dispersal barriers, biotic interactions and local adaptation, all of which interact in complex ways (Martiny *et al.*, 2006). A major endeavour in microbial biogeography has been to disentangle the relative importance of these processes in driving community diversity and turnover (Dumbrell *et al.*, 2010).

Rhizobia are a diverse group of plant symbionts in the α- and β-Proteobacteria. Rhizobia are capable of forming structures known as nodules on the roots and, less frequently, on the stems of most legumes. Within nodules, these bacteria fix atmospheric nitrogen into ammonium that legumes can utilize. In return, legumes provide rhizobia with photosynthates (Denison and Kiers, 2011). Given the tight co-evolved history between legumes and rhizobia, the biogeography of rhizobia is known to be strongly affected by the distribution of compatible host plants, and to a lesser extent by prevailing soil and climatic conditions (Bissett *et al.*, 2010; Lau *et al.*, 2012; Sprent *et al.*, 2017). Importantly, rhizobia can also impact plant distributions by facilitating plant establishment and spread (Traveset and Richardson, 2014), which is a function of the degree of co-evolved specificity between interacting partners. For specificity to evolve, plants need to face nutritional trade-offs that make the investment of photosynthates towards rhizobia worthwhile in terms of increased nitrogen supply and fitness (Denison and Kiers, 2011). Unsurprisingly, legumes have evolved sophisticated ways to reward beneficial strains and sanction non-beneficial, or cheater, rhizobia as a way to counter defection (West *et al.*, 2002; Kiers *et al.*, 2003; Sachs *et al.*, 2010).

In the legume-rhizobium relationship the host plant controls the interaction more strongly when symbiotic benefits are high (Andrews *et al.*, 2017). In conditions where nitrogen is non-limiting, such as those in heavily fertilized cropland, plants relax their defence against defection, consequently allowing for less specific and cooperative rhizobial communities to persist (Kiers *et al.*, 2007; Weese *et al.*, 2015). An apparently similar observation can be made for legumes growing in extremely resource-poor soils and rapidly changing habitats, for example in fire dominated drylands (Hassen *et al.*, 2012). In these habitats, promiscuous plants often interact with a wide diversity of rhizobial symbionts because they cannot exclude rhizobia that might be beneficial when conditions change (i.e. an ecological insurance strategy, Yachi and Loreau, 1999; but see Werner *et al.*, 2018). Examples of this include invasive species such as Australian acacias (Ndlovu *et al.*, 2013; Keet *et al.*, 2017; Le Roux *et al.*, 2018; Dinnage *et al.*, 2019), South African legumes in nutrient-poor soils (Hassen *et al.*, 2012; Lemaire *et al.*, 2015; Le Roux *et al.*, 2017) and legumes in semi-arid regions of Brazil (Radl *et al.*, 2014).

Despite the fact that ecological insurance is often put forward to explain the maintenance of high rhizobial diversity in some promiscuous legume species, manipulative studies often fail to identify a positive relationship between rhizobial diversity and plant nutrition and growth (Bever *et al.*, 2013; Barrett *et al.*, 2015). In nodules colonized by multiple strains, interspecific competition and the presence of cheaters are thought to be responsible for lower plant productivity (Sachs and Simms, 2006; Simonsen *et al.*, 2014; Barrett *et al.*, 2015), rendering the high rhizobial diversity commonly observed in legumes paradoxical (Pahua *et al.*, 2018). The maintenance of high rhizobial diversity has been ascribed to negative density-dependent processes that benefit strains when they are rare (also termed “balancing nodulation”; Siler and Friesen, 2017). This can improve the long-term symbiosis benefits by maintaining rhizobial genetic diversity that may become adaptive in future when environmental conditions change, a strategy that will be particularly relevant in highly dynamic habitats. Indeed, theoretical work has shown that rhizobial diversity can be maintained even in the presence of symbiotically ineffective strains (Bever, 2015).

Given the high context-dependency of plant-mutualist interaction promiscuity, it is maybe unsurprising that agricultural legume production often impacts on the diversity and functioning of symbiotic microbial communities (Kennedy *et al.*, 1995; Hartmann *et al.*, 2015). Tillage, synthetic fertilizer application and monocropping of already-homogenized plant genetic material decreases environmental and host plant diversity that typically underlie symbiotic microbial diversity (Van der Heijden and Wagg, 2012; Vuong *et al.*, 2017). For instance, evidence from agricultural systems, mostly on mycorrhizal fungi, suggests that organic farming practices maintain higher levels of symbiotic microbial diversity than conventional farming (Verbruggen *et al.*, 2010; Hartmann *et al.*, 2015). For rhizobia, such information is scant, partly because a positive link between rhizobial diversity and plant growth has not been established (Barrett *et al.*, 2015). Thus, it remains a mystery whether maintaining rhizobial diversity can have direct positive effects on plant yield in agricultural systems. In line with the ecological insurance hypothesis (Yachi and Loreau, 1999; Siler and Friesen, 2017), it is possible that crops thriving in dynamic environments and under rapidly changing climatic conditions will maintain high levels of rhizobial diversity to ensure symbiotic support over the long run (Ramoneda *et al.*, 2020).

There is increasing awareness that crops need to frequently interact with soil rhizobial propagules in order to maintain the symbiotic fitness benefits derived from these interactions (Burghardt, 2019; Ramoneda *et al.*, 2019). Over their life span, legumes create new available niches for a wide range of rhizobia to colonize (Dinnage *et al.*, 2019), in turn supporting the maintenance of high rhizobial diversity in local plant populations. This may be particularly relevant in perennial legumes under cropping (Burghardt, 2019), making them ideal systems to address how ecological and evolutionary principles can be used to promote beneficial legume-rhizobium interactions. Such legumes allow for the build-up of diverse and site-adapted rhizobial populations that can assist plant growth while buffering their hosts against environmental stresses. Therefore, studying the biogeography of rhizobial symbionts of such crops can inform us on how agricultural practices and environmental and geographical factors define the diversity and structure of root nodule communities (Ramoneda *et al.*, 2020).

Rooibos tea, *Aspalathus linearis* Burm. Dahlgren, provides a unique system to study the biogeography of root nodule communities in both cultivated and wild ancestral populations for numerous reasons. First, this legume is adapted to the extremely nutrient-poor soils of the Core Cape Region of South Africa, a plant biodiversity hotspot undergoing desertification (Malgas *et al.*, 2010; Lötter *et al.*, 2014). Its seeds germinate under elevated soil fertility during post-fire successions, transitioning towards severely nutrient-poor conditions over its life span (Maistry *et al.*, 2015). Second, the species contains distinct ecotypes in the wild (Hawkins *et al.*, 2011), often sympatric with plantations and in a highly heterogeneous landscape where physical barriers to dispersal are common (Cowling *et al.*, 2009; Dludlu *et al.*, 2017). Lastly, recent studies have found rooibos to associate with a wide diversity of rhizobia (Hassen *et al.*, 2012; Le Roux *et al.*, 2017; Ramoneda *et al.*, 2020), particularly at the seedling stage. While the species has a preference for *Mesorhizobium* strains, it has weak control over its rhizobial partners.

In this study we built on the unique ecological features of rooibos to jointly explore the association between agricultural practices, geographical distance, and plant traits with the biogeography and diversity of root nodule communities in association with rooibos. Unlike the majority of studies on microbial biogeography, we used a community prediction approach to *a priori* classify root nodule communities into distinct classes (Arumugam *et al.*, 2011; Pascual-García and Bell, 2019). This method allowed us to objectively describe the structure of the dominant root nodule communities in rooibos, and to relate them to their potential drivers. We specifically asked: 1) Are root nodule communities of rooibos determined by their geographical location?; 2) Do foliar plant traits contribute to explain the structure of such root nodule communities?; and 3) How do conventional and organic rooibos farming impact the composition and diversity of root nodule symbionts? Given the strong habitat heterogeneity and abundant topographical barriers in the landscape, we expected rooibos root nodule communities to be geographically structured (Ramoneda *et al.*, 2020). Moreover, following a similar study describing the root nodule communities of organically cultivated and wild rooibos populations (Le Roux *et al.*, 2017), we expected small compositional and diversity differences between the two. Consequently, we expected geographical distance to be dominant and to have the same effect on root nodule community dissimilarities among cultivated and wild rooibos populations.

## Materials and Methods

### Sample collection

Rooibos is a legume shrub endemic to the Sandstone Fynbos vegetation of the Northern and Western Cape provinces of South Africa. We sampled sites where cultivated and wild rooibos populations occurred within the same area (∼1km^2^) in four main rooibos farming areas: the Suid Bokkeveld, Cederberg, Skimmelberg and Citrusdal (Supp. Fig. 1). Sampling was done in July 2016 (Suid Bokkeveld) and April 2017 (Cederberg, Citrusdal and Skimmelberg). From these sampling areas, we sampled plants and soils from different farms in the Suid Bokkeveld, while in the rest of areas we sampled single but larger farms.

Within each farm, we collected up to ∼ 50 root nodules and > 5 g of leaves from at least seven plants in paired cultivated and adjacent wild rooibos populations. At each site, the seven plants sampled were separated maximally by 50m. When possible, several locations within a farm were sampled. We obtained the root nodules from fine roots in the top 50 cm of soil, and when possible also from lateral roots that extended up to 1 m horizontally from the tap root. After temporary storage in paper bags, root nodules were washed by thoroughly rinsing with pressurized water on the same day and stored them in silica gel to remove all humidity. Rooibos leaves were stored and air-dried in paper bags until further analyses. Within the Suid Bokkeveld, we collected soils from up to 30 cm soil depth in cultivated and adjacent wild populations at five different farms, which were maximally 33 km apart from each other.

Root nodules were weighed and dried leaves were finely ground for nutrient analyses in wolfram carbide cups on a TissueLyser II swing mill (Qiagen, Hombrechtikon, Switzerland) for 60 s at 28 Hz. Root nodules from each individual plant were pooled and then milled using sterile glass beads using a TissueLyser II swing mill for 2 minutes at 15 Hz (Qiagen, Hombrechtikon, Switzerland).

### Analysis of the mineral nutrient concentrations and ^15^N and ^13^C signatures in rooibos leaves and soils

Total foliar nitrogen (N), ^13^C and ^15^N stable isotopes were measured from 4 mg of milled leaf material in tin capsules fed into an NCS analyser (FlashEA 1112 Series, Thermo Fisher Scientific Inc.,Waltham, USA). For phosphorus (P) and other macronutrient measurements, 200 mg of ground leaf material were initially digested in 2 ml HNO_3_ (69%) and 2 ml deionized H_2_O in an MLS Turbowave (MWS GmbH, Heerbrugg, Switzerland). Extracts were then fed into an ICP-OES (ICPE-9800, Shimadzu, Japan). All plant nutrient analyses were carried out in the laboratory facilities of the Group of Plant Nutrition at ETH Zurich (Eschikon, Switzerland). Soil pH was measured in triplicates in a 0.01M CaCl_2_ solution using a pH-meter. The macronutrients and ^15^N isotope measurements in air-dried soil samples were also measured using ICP-OES and NCS elemental analyzer. Specifically, soil extracts were obtained by wet digestion in a mix of 200 mg dry soil, 4 ml of HNO_3_ (69%) and 10ml deionized H_2_O in an MLS Turbowave (MWS GmbH, Heerbrugg, Switzerland).

### DNA extraction, gene amplification and sequencing

DNA was extracted from at least 15 mg of root nodule powder using the NucleoSpin Plant II Kit (Macherey-Nagel, Germany) for each sample following the manufacturer’s protocol. The *gyrB* gene was amplified using the primers *gyrB*343F (5’-GCAGTCGAACATGTAGCTGACTCAGGTCACTTCGACCAGAAYTCCTAYAAGG-3’) and *gyrB*1043 (5’-TGGATCACTTGTGCAAGCATCACATCGTAGAGCTTGTCCTTSGTCTGCG-3’) (Martens *et al.*, 2008), with an amino-5’ end blocking for PacBio sequencing. The *gyrB* gene is a taxonomic marker and housekeeping gene that encodes the B subunit of a gyrase protein, a topoisomerase that introduces negative supercoils to the DNA (Gellert *et al.*, 1979). PCR reactions were performed in triplicates using a final reaction volume of 12.5 µl per sample. Each reaction contained 4.25 µl PCR-grade water, 6.25 µl Multiplex PCR Master Mix (QIAGEN Multiplex PCR Plus Kit), 0.5 µl of each of the primers and 1 µl of the template DNA. Amplification was done in a DNA Engine Peltier Thermal Cycler (Bio-Rad, USA) using the following conditions: an initial denaturation at 95°C for 5 min followed by 30 cycles of 95°C for 30 s, at 58°C for 2 min and 72°C for 45 s, followed by a final extension at 68°C for 10 min. Reactions were kept at 10°C until storage at −20°C. Triplicate PCR reactions were pooled and the amplicons were subsequently purified with self-made SPRI beads at a ratio of 0.7 times the reaction volume (Genetic Diversity Center, ETH Zürich). As a next step the Barcoded Universal F/R Primers Plate - 96 (Pacific Biosciences, USA) with 96 unique barcodes were used to multiplex amplicons in a second PCR step. These barcodes contained a complementary 16-mer sequence to the universal one ligated to the taxonomic primers. The PCR reaction mix contained 12.25 µl PCR-grade water, 5 µl 5x Q5 reaction buffer (New England Biolabs), 2.5 µl dNTPs (2mM), 2.5 µl of each of the index primers, 0.25 µl of Q5 polymerase (New England Biolabs), and 2.5 µl of the template (1 ng µl^-1^). The following PCR cycle was used: 98°C for 30 s, 10 cycles at 98°C for 10 s, 71°C for 20 s and 72°C for 60 s, with a final extension after the cycling at 72°C for 2 min. Products of index PCRs were again purified with the self-made SPRI beads at the same 0.7 sample-to-bead ratio (Genetic Diversity Center, ETH Zürich) and the DNA concentration was recorded with the Qubit assay (Thermo Fisher Scientific Inc., USA) on the Spark 10M Multimode Plate Reader (Tecan, Switzerland).

Size (∼817 bp) and purity of amplicons were determined with in Agilent 2200 Tape Station (Agilent, USA). A total of 316 libraries were finally normalized to the same amount of DNA per sample (10 ng) and pooled to a final concentration of 5 nM. Libraries were sequenced with a PacBio Sequel platform at the Functional Genomics Center Zurich (University of Zurich, Zurich, Switzerland) using the SMRTbell™ Template Prep Kit 1.0-SPv3 (Pacific Biosciences) chemistry for SMRT library preparation. Samples had to be run in 3 different SMRT cells, which generated 3247 sequences per sample on average. An average of 37 polymerase passes per sample, with a minimum of 5, ensured a minimum predicted sequencing accuracy of 0.999.

### Rooibos genetic diversty

Rooibos genetic diversity was characterized by Sanger sequencing of the *psbA-trnH* gene. First, we extracted genomic DNA from 20 mg of milled leaf material using the NucleoSpin Plant II Kit (Macherey-Nagel, Germany). Then, the *psbA-trnH* gene (∼650 bp) was amplified using the primers *psbA3f* (5’-GTTATGCATGAACGTAATGCTC-3’) and *trnHf* (5’-CGCGCATGGTGGATTCACAATCC-3’) (Lahaye *et al.*, 2008). The *psbA-trnH* gene is a chloroplast region that encodes the D1 protein of the photsystem II (Liere *et al.*, 1995), and its low variation is assumed to underpin high genetic divergence even between species. PCR reactions were performed in triplicates using a final reaction volume of 25 µl per sample. Each reaction contained 10.25 µl PCR-grade water, 2.5 µl 10x reaction buffer B (New England Biolabs), 2.5 µl dNTPs (2 mM), 2.5 µl MgCl_2_ (25 mM), 2.5 µl of each of the primers, 0.25 µl FIREPol DNA polymerase (Soils BioDyne, Tartu, Estonia) and 2 µl of the template DNA. Amplification was done in a DNA Engine Peltier Thermal Cycler (Bio-Rad, USA) using the following conditions: an initial denaturation at 95°C for 5 min followed by 22 cycles of 94°C for 30 s, 52°C for 15 s and 72°C for 90 s, followed by a final extension at 72°C for 15 min. Reactions were kept at 10°C until storage at −20°C. Triplicate PCR reactions were pooled and the amplicons were subsequently purified with illustra™ ExoStar™ 1-step (GE. Healthcare, Opfikon, Switzerland). Sequencing premixes for the psbA3f and trnH primers was added to 3 µl of the purified product, containing 2.5 µl PCR-grade water, 2 µl primer (1 µM), 1.5 µl 5x sequencing buffer (New England Biolabs), and 1 µl BigDye (ThermoFisher Scientific, Reinach, Switzerland). Sanger sequencing was performed in an ABI3730 DNA analyzer with 48 capillaries (ThermoFisher Scientific, Reinach, Switzerland) located at the Genetic Diversity Center (GDC) of ETH Zurich.

### Bioinformatics of sequencing data

For the *GyrB* region removal of bell adaptors, barcodes and primer sites, along with demultiplexing, was done using the SMRT link software (Pacific Biosciences). Sequences were filtered by sizes between 700-1000 bp using the same software, resulting in an average fragment length of 801 bp. Quality filtering was done with PRINSEQ-lite (Schmieder and Edwards, 2011). Unique sequences needed for subsequent clustering and annotation were obtained with the USEARCH function *fastx_uniques*. These amplicons were then error-corrected using UNOISE3, which generated ZOTUs (zero-radius operationally taxonomic units). These are expected to cover all unique true biological sequences in the samples, in order to reach a resolution that can distinguish between closely related strains. The ZOTUs were clustered at 97%, 98% and 99% similarity levels with the USEARCH function *cluster_smallmem*, following the recommendations of Edgar (2016). Briefly, provided the UNOISE3 error correction already removed erroneous reads due to sequencing and PCR errors and chimeras, by clustering the reads at 99% similarity we could recover real biological variation. However, we found that clustering at 98% and 99% similarity led to overly high variation that obscured beta diversity patterns. We thus decided to use 97% as the similarity clustering threshold, which had 160 ZOTUs compared to the 201 and 272 ZOTUs of the dataset at 98% and 99% clustering respectively. In this way we were able to characterize the root nodule communities of rooibos despite strain-level variation being high. The count table was obtained with the *otutab* function also in USEARCH. Taxonomic annotation and chimera checking were done with SINTAX (Edgar, 2016), on all available *gyrB* sequences from NCBI GenBank database (https://blast.ncbi.nlm.nih.gov/Blast.cgi).

A total of 426977 *gyrB* sequences were processed (see results section). All raw sequence data processing was done in the *phyloseq* package (v1.16.2) (McMurdie *et al.*, 2013; R Development Core Team, 2016). We excluded a total of fourteen samples with less than 500 reads. We then calculated α-diversity (richness and Simpson’s diversity index) of root nodule communities from intra-and extrapolated ZOTU read counts to the median read number of 1108 reads, using the *iNEXT* package (v.2.0.19) in R (Hsieh *et al.*, 2019; R Development Core Team, 2016). The rest of community analyses were done on relative abundance-corrected counts after verifying that square-root (Hellinger) correction did not change beta diversity patterns. Sequence processing of the *psbA-trnH* chloroplastic gene marker was done using Geneious (*v9.1.8*), where ends were trimmed, paired, and consensus sequences found for each sample. Consensus was established by the highest phred quality score (set to 20 as a minimum) of the two base calls when these were not matching at 100% pairwise identity.

### Determination of community classes

We took an explorative approach to identify the dominant root nodule assemblages in rooibos by clustering the communities according to their structural similarity. We used the method applied by Arumugam *et al.*, (2011) and Pascual-García and Bell (2019) to describe distinct classes of enterotype and treehole bacterial communities. Briefly, we first determined the distance matrix between all communities in the study using Jensen-Shannon (*D*_*JSD*_) divergence (Endres and Schindelin, 2003). Using the R package *CLUSTER* (v2.1.0, R Development Core Team, 2016) we ran the Partitioning Around Medoids (PAM) function on the *D*_*JSD*_ distance matrix. The function requires an *a priori* maximum number of *k* clusters as input, which we set to between 1 and 15, as we did not expect more clusters in the dataset given only 4 regions were sampled and the high simiarity between communities from cultivated and wild rooibos resportd in other studies (Le Roux *et al.*, 2017). To identfy the optimal number of *k* clusters, we computed the Calinski-Harabasz index (*CH*), which quantifies the quality of a classification. The optimal number of *k* clusters was that with the highest *CH* index, which confirmed that the 1 to 15 range was adequate as there was a steady decrease in the *CH* index towards the highest numbers of clusters. We then performed an indicator species analysis to determine which ZOTUs characterized the predicted community classes using the R package *indicspecies* (v1.7.6) and the function *multipatt* (De Caceres and Legendre, 2019; R Development Core Team, 2016). The frequency of the community classes was plotted on maps of the sampled rooibos distribution range generated in Google Earth (*v7.1.7*).

### Statistical analysis

After visualization of the predicted community classes in a Principal Coordinates Analysis (PCoA) with R function *DUDI.PCO* from package *ade4* (*v1.7-13*, Dray and Dufour, 2007; R Development Core Team, 2016), we performed a Permutational Analysis of Variance (PERMANOVA) on 9999 permutations of a Bray-Curtis dissmilarity matrix (*adonis* function of R package *vegan* v2.5-5; Oksanen *et al.*, 2019; R Development Core Team, 2016). This allowed to assess how much variation in community dissimilarity could be explained by the predicted community classes, and to compare it to the factors *Area/Farm/Location within farm* in the outlined nested design, with rooibos population type and genotype as independent factors. We subsequently tested differences in community structure between farms with the *pairwise.adonis* function and related the Pseudo F values of the model output to the geographical distances in km between them. These were then plotted in a scatterplot using *ggplot2* (*v3.2.0*, Wickham 2016; R Development Core Team, 2016). We then used Chi-squared tests on contingency tables to address the relationship between predicted community classes and geographical origin, rooibos population type, conventional versus organic farming, and rooibos genotype.

We used ANOVA to test the effect of farming (conventional/organic) on the intra-/extrapolated observed species (i.e. richness) and Simpson’s diversity index (D). The same approach was used to evaluate the relationship between plant traits and community classes. Contrasts between the mean plant traits of the different community classes were obtained using post-hoc Tukey’s HSD tests at 95% confidence. We also summarized rooibos trait variation using a Principal Components Analysis (PCA) with all measured foliar traits as input. We then overlaid arrows indicating variation in the traits significantly related to community classes.

The links between soil nutrient concentrations and root nodule community structure were determined using correlational approaches. Since we only had soil data from single farm x rooibos population type (cultivated/wild) combinations, the use of linear models was not possible due to strong autocorrelation. Instead, beta diversity was reduced to distances between centroids in a PCoA ordination based on a Bray-Curtis dissimilarity matrix (the centroid value of all replicates of the farm x rooibos population type combination in the euclidean space), and these values were then correlated to the euclidean distances between soil parameter values. This was done using the *dist_between_centroids* function of the *usedist* package (*v0.1.0*, https://CRAN.R-project.org/package=usedist; R Development Core Team, 2016). In this way correlations between the soil nutrient concentrations and root nodule community structure could be tested without violating fundamental statistical assumptions.

## Results

We sampled 286 rooibos plants and, based on the *gyrB* gene, characterized their respective root nodule communities in paired cultivated and wild populations in the rooibos geographical distribution range. We investigated alpha and beta diversity patterns of the rooibos root nodule communities and related them to the geographical locations, rooibos population types and leaf traits of the plants where they were sampled. Using a statistical community prediction approach, we were able to identify the major root nodule assemblages of rooibos and describe the drivers of their biogeography.

### Rooibos root nodule communities are defined by the dominance of distinct and widespread Mesorhizobium strains

We detected a total of 160 *gyrB* ZOTUs in rooibos root nodules when clustered at 97% similarity. These belonged to the genera *Bradyrhizobium* (69), *Rhizobium* (26), *Mesorhizobium* (7), *Agrobacterium* (3), and *Tardiphaga* (1), while 54 ZOTUs could only be identified at the family level or above. The community prediction approach based on Jensen-Shannon divergence (D_JSD_) identified four distinct *gyrB* root nodule community classes (Fig. 1A; Suppl. Figure 2). These were all dominated by ZOTUs of the genus *Mesorhizobium*, and the communities were strongly differentiated by the dominance of distinct ZOTUs of this genus (Fig. 1B). One of these, ZOTU 420, attained considerable relative abundance (>20 %) in all community classes, and defined community class 3 (Fig. 1B). All other community classes were characterized by the dominance (>50 % relative abundance) of *Mesorhizobium* ZOTUs that were widespread but rare in the other community classes.

**Figure 1.**
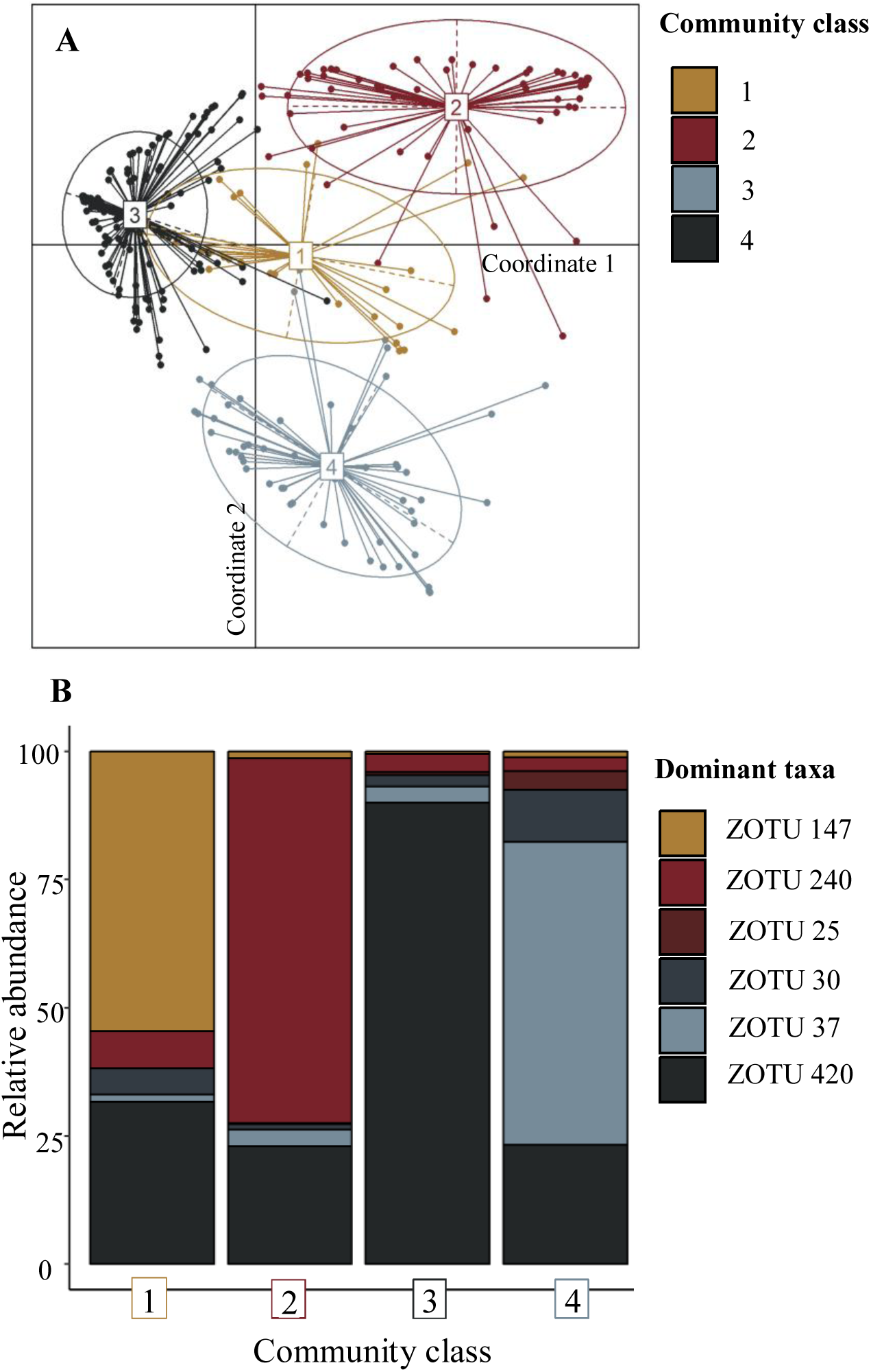
Description of the four root nodule community classes of rooibos based on a partitioning around medoids (PAM) approach. (A) Principal Coordinates Analysis (PCoA) of rooibos root nodule communities based on Jensen-Shannon distances. Different colours depict the four root nodule community classes predicted by the PAM function after *k*-clustering with the Calinski-Harabasz (CH) index. (B) Stacked barplot showing the relative abundance of the six most abundant ZOTUs across all samples (n = 286) in the four predicted community classes. All ZOTUs belonged to the genus *Mesorhizobium*, except for ZOTU 30 (*Rhizobium*). The analysis relied on *gyrB* phylotaxonomic marker sequences and their comparison to a reference database of all publicly available and taxonomically assigned sequences. Ellipses indicate 95% confidence intervals around the centroid.

The indicator species analysis revealed more details about the compositional differences between the community classes. Firstly, community class 3 was the only assemblage where no indicator ZOTUs were obtained (Table 1). Along with the fact that half of the samples clustered within this community class (N = 144), this finding indicates community class 3 is the archetypal root nodule community in rooibos root nodules of this study. Community class 1 (N = 28) was characterized by the dominance of a widespread *Mesorhizobium* ZOTU and a rare *Rhizobium* ZOTU. In contrast, apart from the dominance of a different *Mesorhizobium* ZOTU, community class 2 (N = 59) was characterized by a set of 12 site-specific *Bradyrhizobium* ZOTUs endemic to the Skimmelberg mountain range. Finally, community class 4 (N = 55) had, apart from a dominant *Mesorhizobium* ZOTU, 4 additional rare *Bradyrhizobium* and *Rhizobium* ZOTUS (Table 1).

**Table 1.**
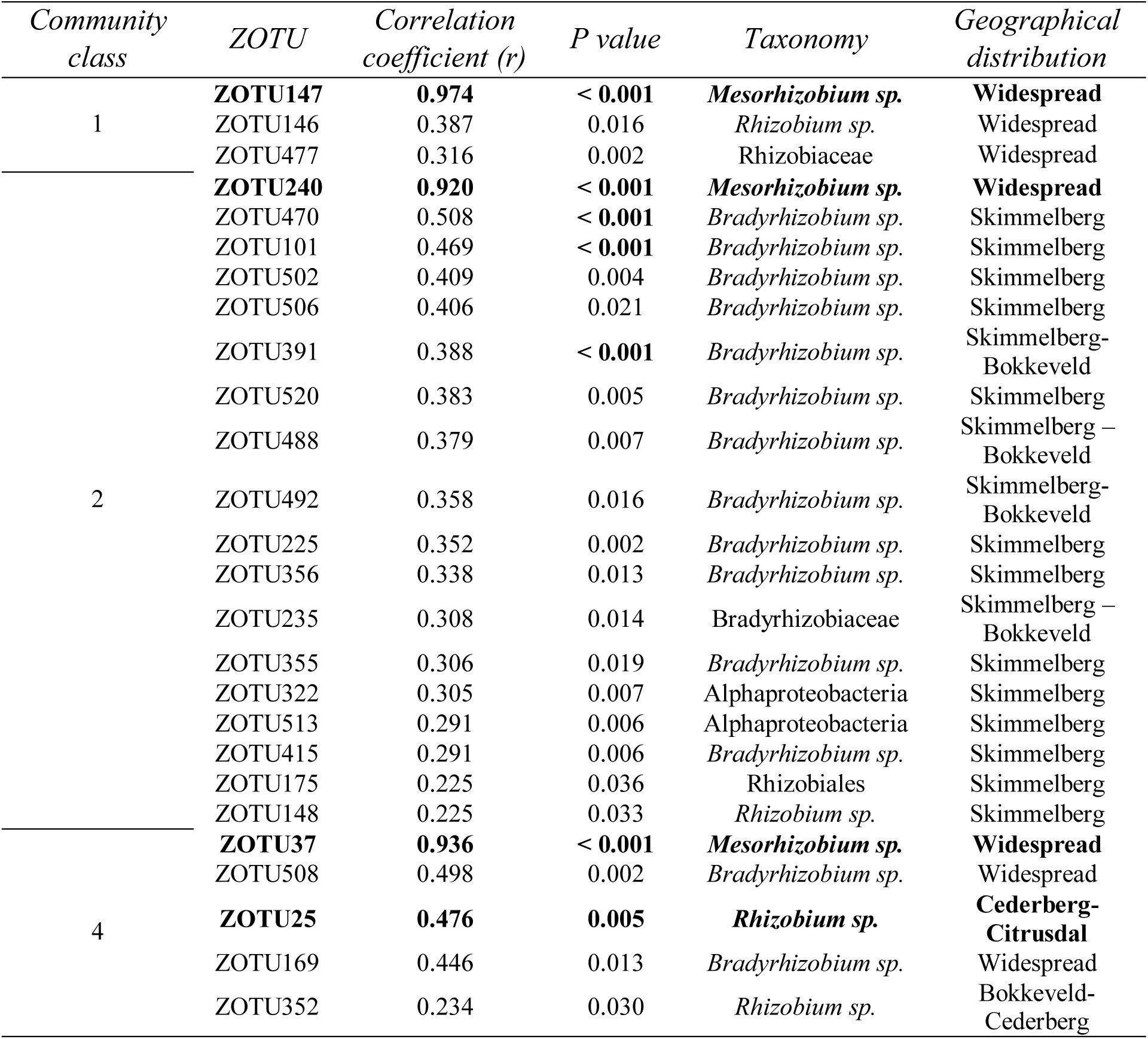
Indicator species analysis of the community classes identified in rooibos root nodules using a partitioning around medoids (PAM) approach. Only significant (P < 0.05) correlations are shown and taxa were classified according to their distribution range, which was widespread when they occurred at least once in every farming area. Community class 3 was the only assemblage where no indicator ZOTUs were obtained.

### Root nodule community classes are associated with rooibos geography and population type (planted and wild), but not with plant genotype

We addressed whether rooibos root nodule community structure was influenced by geographic location, rooibos population type (planted versus wild) or rooibos genotype. Pearson’s Chi-squared tests found significant relationships between the geographic origin and rooibos population type of these communities, but not with rooibos genotype (Fig. 2). The strength of the association between community classes and geographical locations increased with spatial resolution in the sequence Geographical Area < Farm < Locality within farm (Table 2).

**Table 2.** Chi-squared tests of relationships between predicted root nodule community classes and geographical locations, rooibos population type (cultivated vs wild), farming practice (conventional vs organic) and genotype, across the studied rooibos distribution range. Rooibos genotypes were determined after aligning a 650bp fragment of the *psbA-trnH* chloroplastic marker from plants in which the root nodule communities were described.

| Variable | $\chi^2$ statistic | df | P-value |
| --- | --- | --- | --- |
| Area | 91.49 | 63 | < **0.001** |
| Farm | 138.47 | 27 | < **0.001** |
| Farm locality | 202.65 | 9 | < **0.001** |
| Population type | 19.55 | 3 | < **0.001** |
| Farming practice | 8.85 | 3 | **0.031** |
| Genotype | 19.52 | 21 | 0.552 |

**Figure 2.**
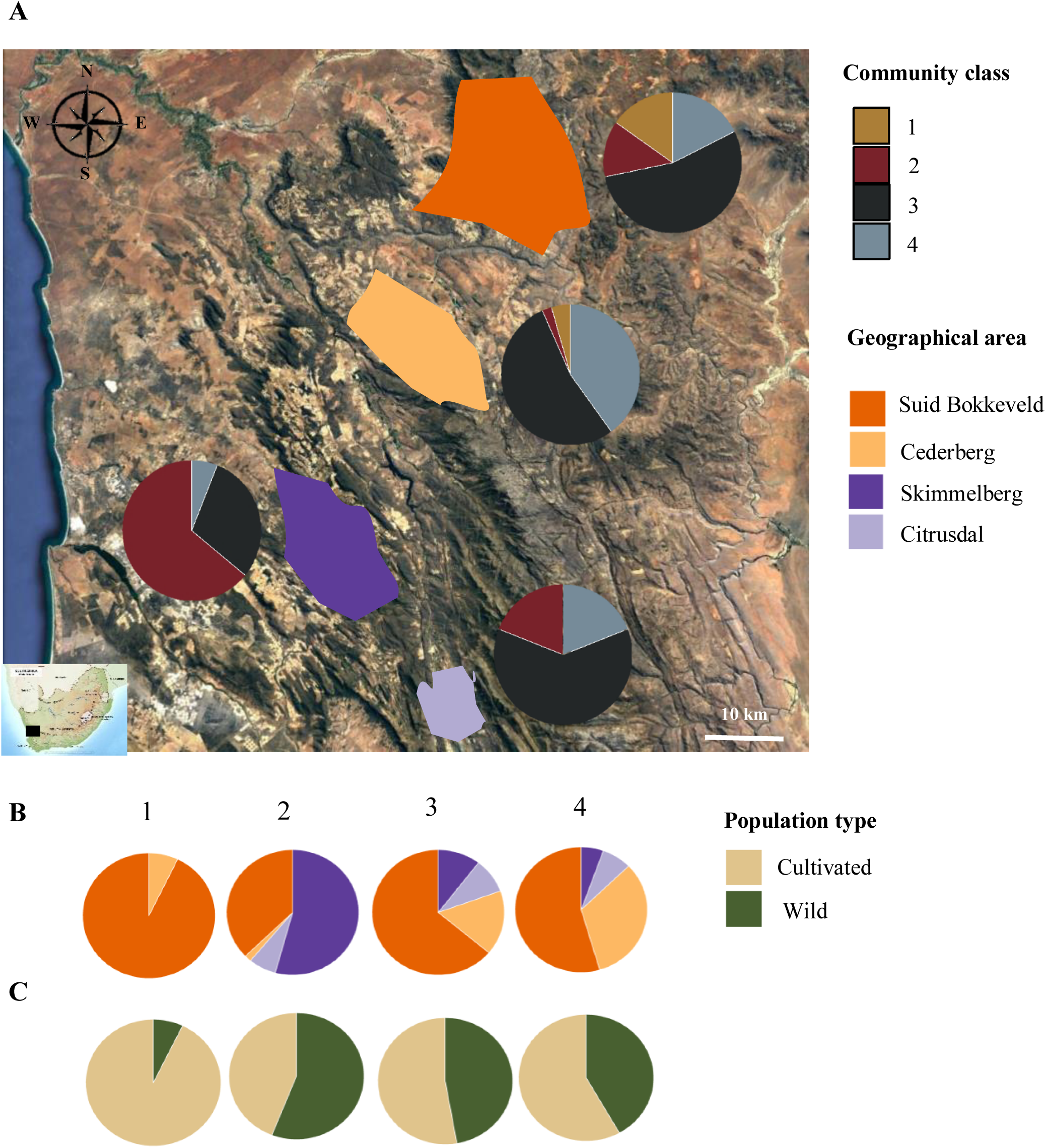
Distribution of the root nodule community classes in the areas sampled in this study. (A) Map of the rooibos cultivation areas sampled in this study with the proportion of samples of each of the four root nodule community classes identified found in each area. (B) Proportion of samples from each of the rooibos cultivation areas belonging to each of the four root nodule community classes identified (numbers 1 to 4). (C) Proportion of samples from each of the four root nodule community classes identified found in either cultivated or wild rooibos populations.

Although rooibos population type had only a minor association to community classes, in the Skimmelberg area wild plants contained a majority of distinctive *Bradyrhizobium* strains of community class 2 (Fig. 2). This class was the only one with more communities from wild than cultivated rooibos populations. This is in contrast to community class 1, which was characteristic of a number of plants from rooibos plantations in the Suid Bokkeveld (Fig. 2).

Finally, community class 4 was most characteristic of cultivated and wild rooibos plants from the Cederberg mountain range (farm Klipopmekaar, Fig. 2). Up to 40% of the communities sampled in this area belonged to this community class, with the rest of Cederberg communities being clustered almost entirely in the archetypal community 3 (Fig. 2).

### Root nodule beta diversity turnover

The link between community classes and geographical locations and population types of rooibos suggests that these attributes play a role in root nodule community structure. We quantified the variation explained by geographical scale and population type using PERMANOVA with a nested design (see Materials and Methods), followed by a pairwise PERMANOVA to quantify differences between factor levels.

The predicted community classes explained most of the variation in Bray-Curtis dissimilarity between rooibos root nodule assemblages (R^2^ = 0.64, P < 0.001; Table 3). Geographical factors such as the region, farm and location within farm, together only explained 32.1% of the variation in root nodule community structure (Table 3). The effect of rooibos population type alone was weaker (R^2^ = 0.02, P = 0.005), and importantly, there was no effect of rooibos genotype (R^2^ = 0.01, P = 0.543, based on N_genotype A_ = 104 and N_genotype G_ = 25). Finally, conventional rooibos farming had a very marginal impact on root nodule community structure compared to organic farming (R^2^ = 0.01, P = 0.069).

**Table 3.** Summary of the Permutational Analysis of Variance (PERMANOVA) based on Bray-Curtis dissimilarities of rooibos root nodule communities across the plant’s geographical distribution range covered in this study. The variable ‘community type’ corresponds to the predicted root nodules community classes of rooibos using a partitioning around medoids (PAM) approach. The factor ‘Farm’ was nested within ‘Area’ and ‘Location within farm’ was nested within ‘Farm’.

| Variable | df | F-value | $R^2$ | P-value | Unexplained variation (%) |
| --- | --- | --- | --- | --- | --- |
| Community type | 3 | 164.20 | 0.636 | < **0.001** | 36.4 |
| Area | 3 | 7.12 | 0.068 | < **0.001** |  |
| Area : Farm | 6 | 2.54 | 0.049 | < **0.001** | 77.9 |
| Area : Farm : Location within farm | 12 | 3.27 | 0.104 | < **0.001** |  |
| Population type | 1 | 4.26 | 0.015 | **0.005** | 98.5 |
| Farming | 1 | 2.22 | 0.008 | 0.069 | 99.2 |
| Genotype | 1 | 0.76 | 0.006 | 0.543 | 99.4 |

### Rooibos farming impacts the alpha and beta diversity of root nodule communities differently

Characterization of root nodule rhizobial richness (observed species) showed significantly lower richness in conventional (mean ± SD; obs. species = 18.12 ± 10.64, N = 27) than organic (obs. species = 27.28 ± 12.92, N = 120; P < 0.001) rooibos plantations (Fig. 3A). The same differences were observed for Simpson’s D (Fig. 3B). Additionally, richness and diversity were higher in organically farmed rooibos (obs. species = 27.28 ± 12.92, D = 3.19 ± 1.58, N = 120) than in wild populations (obs. species = 22.92 ± 12.19, D = 2.58 ± 1.38, N = 103; P = 0.01).

**Figure 3.**
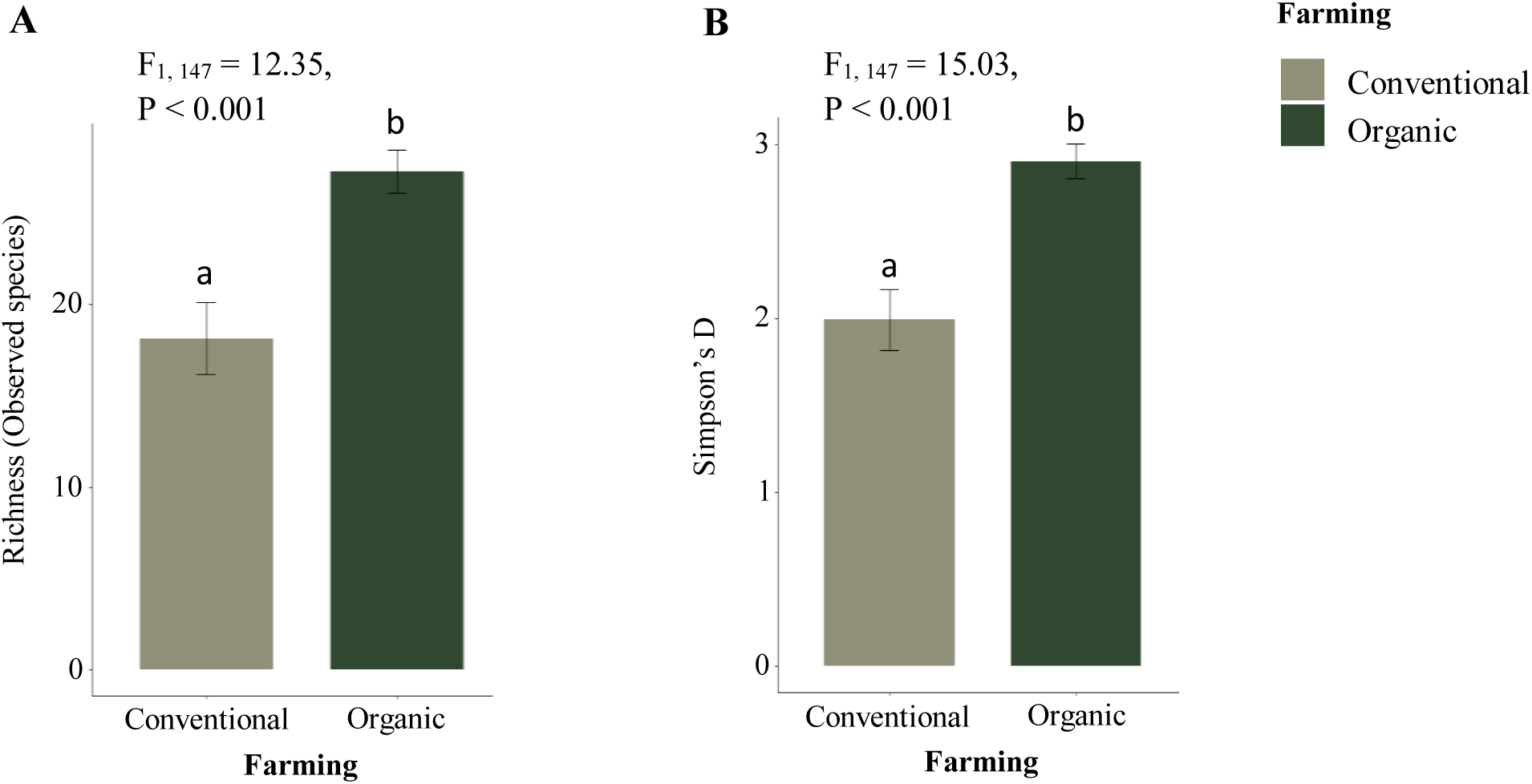
Effects of conventional versus organic farming practices on the diversity of root nodule communities of rooibos. The ZOTUs were determined by similarity clustering of a 817 bp fragment of the phylotaxonomic marker gene *gyrB* at 97% sequence identity. The statistical summary reports differences in richness and Simpson’s D between conventionally and organically farmed rooibos (n=147). Error bars depict ±2 standard errors (SE) around the mean.

The substantial effects of geographical location on root nodule community structure opened the question of whether these effects became stronger with increasing distance between sites. We found a distance-decay pattern of community similarity in which geographical distance (km) correlated positively with community dissimilarity (Bray-Curtis dissimilarity) (Fig. 4). Interestingly, this pattern was only present in wild rooibos populations (r = 0.572, P < 0.001; Fig 4A), whereas no decay in community similarity with increasing distance was observed in plantations (r = 0.184, P = 0.284; Fig. 4B).

**Figure 4.**
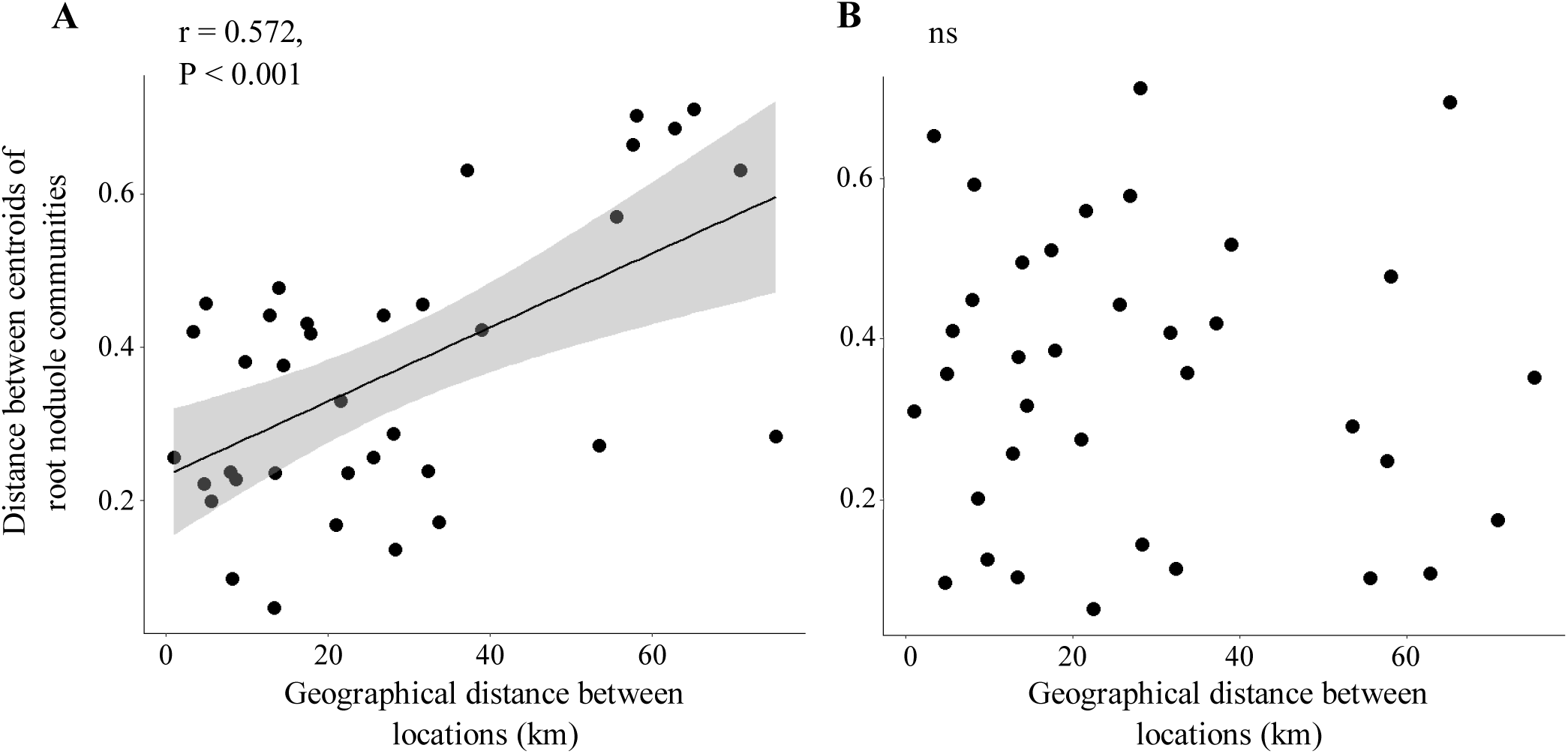
Linear regressions of the relationship between geographical distance and root nodule community dissimilarity based on *gyrB* sequencing data. (A) Significant distance-decay relationship of community similarity from root nodule communities obtained from wild rooibos populations. (B) Absence of a distance-decay relationship of community similarity between root nodule communities obtained from cultivated rooibos populations. Dots depict distances between centroids of Bray-Curtis dissimilarity values between root nodule communities.

### Links between community classes and rooibos foliar phenotypic traits

We used linear models and PCA to test whether rhizobial community classes were associated with particular rooibos foliar nutrient stoichiometric traits. From all measured foliar nutrient concentrations and stable isotopes, we found significant associations between rhizobial community types and foliar ^15^N (‰), ^13^C (‰), N (mg g^-1^), C:N ratio, Mg (mg g^-1^) and Mn (mg g^-1^) (Table 4, Fig. 5A-F). However, a PCA ordination combining all measured traits shows that there was no integrated phenotype associated with community classes (Fig. 6). Instead, these seemed to be associated with discrete traits. The analysis of rooibos genotypic variation only identified two dominant genotypes (Supp. Fig. 3). In line with the finding that community classes were rather related to discrete foliar traits, these genotypes were not associated with any particular community class. This is so despite rooibos genotypes A and G showed signs of phenotypic variation (Supp. Fig 3).

**Table 4.** Relationships between foliar stoichiometric traits of rooibos and predicted root nodule community classes tested with a generalized linear model. Rhizobial community classes were identified using a partitioning around medoinds (PAM) function. Statistical significance was set at P ≤ 0.05.

| Plant traits | N | Community class (df = 3) |  |  |  | Model output |  |  |
| --- | --- | --- | --- | --- | --- | --- | --- | --- |
|  |  | 1 | 2 | 3 | 4 | F-value | R<sup>2</sup> | P-value |
| Foliar <sup>15</sup>N (‰) | 262 | 1.52 ± 1.79 | 0.32 ± 1.36 | 0.88 ± 1.53 | 1.36 ± 2.16 | 4.73 | 0.04 | **0.003** |
| Foliar <sup>13</sup>C (‰) | 221 | -29.97 ± 1.62 | -30.31 ± 2.05 | -29.60 ± 2.13 | -29.05 ± 2.10 | 2.87 | 0.02 | **0.040** |
| Foliar N (mg g<sup>-1</sup>) | 262 | 16.61 ± 1.85 | 14.13 ± 2.43 | 15.22 ± 2.39 | 15.48 ± 2.51 | 6.93 | 0.06 | **< 0.001** |
| Foliar C:N ratio | 262 | 29.66 ± 3.92 | 35.44 ± 5.92 | 33.01 ± 5.51 | 32.33 ± 5.94 | 6.84 | 0.06 | **< 0.001** |
| Foliar P (mg g<sup>-1</sup>) | 249 | 0.50 ± 0.08 | 0.48 ± 0.19 | 0.47 ± 0.20 | 0.49 ± 0.17 | 0.38 | 0.00 | 0.766 |
| Foliar N:P ratio | 249 | 33.47 ± 5.63 | 32.77 ± 12.21 | 36.88 ± 12.55 | 33.38 ± 10.62 | 2.19 | 0.01 | 0.09 |
| Foliar Ca (mg g<sup>-1</sup>) | 257 | 1.78 ± 0.92 | 2.43 ± 1.15 | 2.20 ± 1.00 | 2.26 ± 1.19 | 2.10 | 0.01 | 0.101 |
| Foliar K (mg g<sup>-1</sup>) | 247 | 3.58 ± 0.87 | 3.59 ± 1.17 | 3.83 ± 1.09 | 3.85 ± 1.27 | 0.83 | 0.01 | 0.476 |
| Foliar Mg (mg g<sup>-1</sup>) | 255 | 1.68 ± 0.50 | 2.89 ± 1.06 | 2.49 ± 0.88 | 2.38 ± 0.93 | 10.31 | 0.10 | **< 0.001** |
| Foliar Mn (mg g<sup>-1</sup>) | 250 | 0.11 ± 0.05 | 0.15 ± 0.10 | 0.11 ± 0.07 | 0.16 ± 0.23 | 3.29 | 0.03 | **0.020** |
| Foliar Fe (mg g<sup>-1</sup>) | 255 | 0.09 ± 0.05 | 0.09 ± 0.05 | 0.09 ± 0.09 | 0.08 ± 0.04 | 0.11 | 0.00 | 0.957 |

Community classes 1 and 2 were most dissimilar in their overall foliar stoichiometric traits, particularly in foliar ^15^N, C:N ratio, and Mg concentration. Since these community classes were preferentially found in cultivated and wild rooibos populations respectively, we tested how much variation in these stoichiometric traits could be explained by population type, i.e. plantation vs. wild populations. Indeed, foliar ^15^N and C:N ratios were most explained by population type (^15^N: R^2^ = 0.266, P < 0.001; C:N ratio: R^2^ = 0.119, P < 0.001), with community classes explaining much smaller amounts of variation (^15^N: R^2^ = 0.041, P = 0.003; C:N ratio: R^2^ = 0.062, P < 0.001) (Table 4). Instead, the strongest explanatory variable for foliar Mg concentration was farm location (R^2^ = 0.158, P < 0.001), followed by area (R^2^ = 0.100, P < 0.001), community class (R^2^ = 0.097, P < 0.001) and population type (R^2^ = 0.067, P < 0.001). This seems to be related to the preferential presence of community class 2 in the Skimmelberg farm (Fig. 3).

## Discussion

In this study we analysed the geographical distribution and structure of root nodule rhizobial communities in association with rooibos in cultivated and wild populations throughout the species range. Using a community prediction approach, we show that rooibos contains distinct root nodule community classes strongly linked to geographical location, plant leaf traits, rooibos population types (wild versus planted) and soil nutrient concentrations. Moreover, we found that rooibos farming influences rooibos root nodule communities by i) decreasing taxonomic diversity under conventional but not organic farming, and ii) homogenizing root nodule community composition in the distribution range of the plant. We therefore reject our main hypothesis that geographical distance would have the same effect on the dissimilarity of rhizobial communities found in cultivated and wild rooibos populations. Below we discuss the complex combination of soil conditions, plant traits, and dispersal history in driving the biogeography of rooibos root nodule symbionts.

Rooibos root nodule communities could be classified into four main classes. These were defined by the dominance of different *Mesorhizobium* taxa that had widespread geographical distributions but became locally abundant. Moreover, a number of rare *Bradyrhizobium* taxa were also characteristic of particular community types, in particular those in the Skimmelberg mountain range. The role of *Mesorhizobium* as the core rhizobial symbionts of rooibos has been reported a number of times before (Hassen *et al.*, 2012; Lemaire *et al.*, 2015; Le Roux *et al.*, 2017; Ramoneda *et al.*, 2020), and the genus is considered generalist in terms of soil environmental tolerance, particularly towards pH (Dludlu *et al.*, 2018). While edaphic factors (e.g. nutrient or water availability) often explain some variability in root nodule community structure (Vuong *et al.*, 2017), the host plant is normally the strongest filter to rhizobial propagation during their endophytic life stage (Denison and Kiers, 2011). Indeed, legume composition in the Core Cape Region is known to be strongly driven by edaphic factors (Dludlu *et al.*, 2017), suggesting the effects of edaphic factors on root nodule communities are most likely mediated by plant hosts.

Over larger spatial scales, rhizobial communities may differ due to different vegetation types favouring different rhizobium populations in different areas (Yan *et al.*, 2014; Sprent *et al.*, 2017; Pahua *et al.*, 2018). In a similar manner, intraspecific genetic variation is known to affect the stability of legume-rhizobium associations by regulating the specificity of the interaction (e.g. in *Medicago sp.*, Simonsen and Stinchcombe, 2014; Wendtland *et al.*, 2019). Ecotypic variation in rooibos may therefore be an additional filter to root nodule community structure (Malgas *et al.*, 2010; Hawkins *et al.*, 2011), but remains to be tested. In this study we did not find any links between rooibos genotype and root nodule community structure, despite the low genetic variation captured in our chloroplast marker (Supp. Fig. 3).

We found a significant association between community classes and foliar plant traits, which suggests that i) rooibos has distinct functional responses to different root nodule assemblages (e.g. particular assemblages being more efficient at fixing N_2_ than others), or ii) different rhizobia have preference for rooibos populations with particular leaf traits, which can reflect both the physiological and health status and ecotypic identity of rooibos. While both possibilities are not mutually exclusive and have been reported in other studies (Heath, 2010; Van Cauwenberghe *et al.*, 2016; Keller and Lau, 2018), inferences about rhizobial functioning in this field survey can only be speculative. A myriad of unrecorded environmental factors, from soil depth and vegetation cover to climatic ones, are known to affect leaf stoichiometric traits in different ways (Stewart *et al.*, 1995). For example, plants in the North of the Suid Bokkeveld may receive up to twice as much rainfall as those in the lower Cederberg range (Hawkins *et al.*, 2011), rainfall being one of the many factors known to impact both foliar macronutrient concentrations and isotopic signatures (Stewart *et al.*, 1995). Despite these caveats, the relationships between leaf stoichiometric traits and root nodule community types is an interesting finding for a promiscuous plant like rooibos, and deserves further attention in future studies.

Among all possible drivers of root nodule community structure, geographical location and distance were the most prominent ones. While at a local scale (i.e. at the farm and within farm level) the environmental factors discussed above can play a complex but dominant role (Yan *et al.*, 2014; Vuong *et al.*, 2017), at regional scales (i.e. at the area level, in this study at distances of 30-100 km) the drivers are not related to the environmental conditions (Hanson *et al.*, 2012). Instead, the distinct assemblages of rhizobia found in the four different areas reflect a degree of isolation driven by barriers to dispersal, such as the wide valleys and ridges that separate the four areas studied here (Martiny *et al.*, 2006; Dumbrell, 2010; Zhou and Ning, 2017; Suppl. Fig. 1).

Dispersal in rhizobia has received little attention (Madsen and Alexander, 1982; Van Cauwenberghe *et al.*, 2015), although it is considered to be active at local scales and passive at regional scales. Passive dispersal implies all strains are equally able to disperse to other areas (i.e. neutral dispersal). Such neutral dispersal offers opportunities for microbial communities to become differentiated and diverge through ecological and genetic drift (Martiny *et al.*, 2006; Hanson *et al.*, 2012; Zhou and Ning, 2017). This may explain the widespread nature of all dominant *Mesorhizobium* ZOTUs, which might be subject to changes in their abundances due to ecological drift when in isolation (Hanson *et al.*, 2012). The strength of ecological drift is indeed assumed to be stronger in functionally redundant taxa such as the case of *Mesorhizobium* in rooibos (Hassen *et al.*, 2012; Zhou and Ning, 2017). The fact that we find a number of endemic *Bradyrhizobium* taxa in Skimmelberg is also suggestive that rhizobia in rooibos can become isolated, particularly in such a heterogeneous landscape (Cowling *et al.*, 2009; Beukes *et al.*, 2016; Dudlu *et al.*, 2018; Stepkowski *et al.*, 2018). However, even in Skimmelberg or the Cederberg mountain ranges we identified all rhizobial community types, indicating that soil rhizobial populations still underpin soil legacies from historical events of dispersal and changing habitat conditions (Stepkowski *et al.*, 2018). Historical soil legacies can thus explain why both endemism and generalism are represented in rooibos root nodule communities.

The key finding of a positive distance-decay relationship of community similarity in wild rooibos populations is another strong indicator that regional factors drive rooibos root nodule communities through partial geographical isolation (Hanson *et al.*, 2012). Such patterns have been reported in other bacterial groups, but rarely in rhizobia (Horner-Devine *et al.*, 2004; Zhao *et al.*, 2014; Stefan *et al.*, 2018). Given the known specificity of rhizobia for particular soils and plant species, however, these patterns may be widespread in heterogeneous landscapes such as the Core Cape Floristic Region (Stepkowski *et al.*, 2018). The general distance-decay pattern of community similarity was also assessed within the Suid Bokkeveld, which contained farms maximally 32 km apart, as a way to reveal the spatial scale at which root nodule communities became differentiated. We found no relationship between community dissimilarity and geographical distance within the Suid Bokkeveld (Supp. Fig. 5), in agreement with a recent study from the same area (Ramoneda *et al.*, 2020). However, Ramoneda *et al.*, (2020) found a distance-decay pattern only when a functional marker (*nodA*) was considered. We then tested if the pattern held for the present study using the same *nodA* gene, but again found no correlation (Supp. Fig. 6).

Taken together, our results have several implications for our understanding of root nodule community assembly in plants adapted to extreme nutrient limitation and dynamic environments like rooibos. Firstly, root nodule communities become differentiated only when large barriers such as mountain ranges and valleys limit passive rhizobial dispersal (e.g. farms in the Suid Bokkeveld lie on a plateau, whereas Skimmelberg and Cederberg are mountain ranges separated by wide valleys, Citrusdal being at the bottom of one of the valleys). Secondly, the life stage of rooibos has a strong effect on its root nodule community structure. This is so because Ramoneda *et al.*, (2020) reported that the distance-decay pattern of community similarity was observed in evenly aged, 8-month old rooibos seedlings, whereas the lack of pattern found here is among plants between 1.5 and approximately 15 years old. Thus, there seems to be a filtering of rooibos root nodule symbionts during growth towards its mature stages, leading to higher community similarity between sites and a stronger dominance of the genus *Mesorhizobium*. Changes in the root nodule symbionts over time have been reported in other legumes (e.g. in acacias, Dinnage *et al.*, 2019), which in a perennial plant like rooibos may respond to the nutritional trade-offs triggered by the strong changes in nutrient availability characteristic of the transition from seedling to maturity (Burghardt, 2019).

While the diverse effects of farming practices on root symbiont diversity are widely known for many crops (Shannon *et al.*, 2002; Verbruggen *et al.*, 2010; Yan *et al.*, 2014; Hartmann *et al.*, 2015), these had never been assessed in rooibos. We show that organic rooibos farming maintains higher levels of rhizobial diversity in the nodules than wild populations, while conventional farming clearly decreases it. Organic rooibos farming is characterized by extremely low external inputs and no tillage and is practised in small fields within extensive wild vegetation patches in its native distribution range (Hawkins *et al.*, 2011; Lötter *et al.*, 2014). In these areas the farmers often leave plant material remaining from previous harvests and even weeds on the cropping ground as a way to maintain soil organic matter and moisture. These conditions resemble wild habitats that have higher vegetation cover, organic matter and moisture. Considering the fact that in the wild rooibos plants grow in isolation, it seems plausible that organic rooibos plantations act as hubs of rhizobial diversity derived from the surrounding wild vegetation that is further maintained by the larger and denser rooibos population sizes that characterize them. Conversely, conventional farming occurring at the margins of the native distribution range, happening in larger fields without neighbouring wild rooibos populations, and under higher external nutrient inputs seems to compromise rhizobium diversity.

The fact that a distance-decay pattern of community similarity was found in wild but not cultivated populations indicates rooibos farming had a homogenizing effect on the root nodule communities. This implies that, despite locally organic rooibos farming maintained higher root nodule diversity, farming practices end up decreasing symbiont diversity at regional scales. Removal of alternative host plants, soil preparation, and growth of a single cultivar reduced differentiation between rooibos root nodule communities, thus removing the effect of distance and isolation (Hanson *et al.*, 2012). Moreover, in plantation rooibos plants are evenly aged, which may further contribute to equalizing the composition of root nodule communities in plantations and not wild populations. Despite we found little evidence for an effect of plant genotype or leaf stoichiometric phenotype (Supp. Fig. 3), in the wild there are many distinct ecotypes (Malgas *et al.*, 2010; Hawkins *et al.*, 2011), whereas plantations are mostly dominated by a single commercial genotype. Differential effects of cropping on alpha and beta diversity are important to consider when assessing the impacts of agricultural practices on root symbiotic diversity.

Within the Suid Bokkeveld, two of the same four community classes identified were associated with distinct soil N and K concentrations. The fact that 94% of root nodule communities in class 1 (the only one with significantly lower average soil N concentrations) belonged to cultivated populations, indicates the association might be confounded by rooibos population type. Moreover, the lack of correlation between community similarity and any of the soil conditions measured, indicates soil abiotic factors are weak drivers of root nodule community structure in rooibos (Suppl. Fig. 5). This is so despite community class 1 being widespread across all farms in the Suid Bokkeveld (Supp. Fig. 7), leading to the overall conclusion that soil abiotic factors *per se* may not have direct effects on rhizobia nodulating rooibos roots. Instead, soil factors may be indirectly influencing root nodule community structure by regulating the composition and physiology of surrounding plant communities and the rooibos physiological status. This idea would be confirmed by the joint assessment of plant and root nodule community composition in a variety of soil conditions across the Core Cape Floristic Region.

## Conclusions

In this study we described ecological patterns of rooibos root nodule communities from cultivated and wild populations using, for the first time, a community prediction approach. We found that despite organic rooibos farming promoting root nodule community diversity locally, agricultural practices contribute to a homogenization of rooibos root nodule communities regionally. This was revealed by the absence of a distance-decay pattern of community similarity in plantations that was present among wild populations at regional scales. This finding, along with the strong association between rooibos farming regions and predicted community classes, indicates barriers to dispersal seem to be dominant drivers of root nodule community structure in rooibos. The evidence that particular plant traits and soil conditions are also associated to root nodule community structure indicate that rooibos ecotypes found under different environments may harbour distinct root nodule communities.

The findings of this study agree with those reported by Ramoneda *et al.*, (2020) who recorded the early assembly of root nodule communities in rooibos. While at the seedling stage plants allow for a wide diversity of symbionts into the roots, over time there seems to be a filtering towards a stronger dominance of *Mesorhizobium* taxa. This may be due to the natural shift towards nutrient-limiting conditions over the plant’s lifespan, which increases the filtering towards better performing strains (Kiers *et al.*, 2003). In the context of mechanisms of root nodule community assembly in rooibos, there are three main conclusions that can be drawn: 1) the lack of symbiont filtering at the seedling stage underpins predominantly neutral (i.e. stochastic) assembly processes in rooibos root nodule community assembly; 2) over time, the assembly becomes increasingly trait-based (i.e. deterministic) through the filtering by the plant as affected by soil, surrounding vegetation and climate; and, 3) at a regional scale barriers to dispersal enforce isolation and ecological drift between rooibos root nodule communities, a neutral process that enhances dissimilarities between them. Finally, human intervention can modulate rooibos root nodule community structure and diversity through manipulation of the plant’s genetic material and habitat modification. This study shows that a deliberate manipulation of the rooibos root microbiome would need to account for both regional and local ecological processes that drive the structure of its root nodule communities.

## Acknowledgements

This study was funded by the Mercator Research Program of the World Food Systems Center of ETH Zurich. Data produced and analyzed in this paper were generated in collaboration with the Functional Genomics Center of the University of Zurich (FGCZ) and the Genetic Diversity Center of ETH Zurich (GDC). JR acknowledges the assistance from Dr. Jean-Claude Walser, Dr. Andrea Patrignani and Dr. Weihong Qi for bioinformatical analyses, to Stefanie Stadelmann and Sandra Wenger for their assistance in the lab and field respectively, and to Dr. Laurie Schönholtzer and Federica Tamburini for the plant and soil nutrient analyses. JLR acknowledges funding from Macquarie University’s Faculty of Science and Engineering and Department of Biological Sciences. Finally, the authors acknowledge the assistance of Noel Oettlé, Dr. Cecilia Bester, and the community of rooibos farmers in the Suid Bokkeveld for making the sampling in South Africa possible. The authors declare no conflict of interest.

