## Supplementary Material for "Ecological patterns of root nodule diversity in cultivated and wild rooibos populations: a community prediction approach"

**Figure S1** Map of the rooibos cultivation areas sampled in this study.

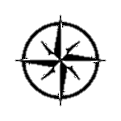

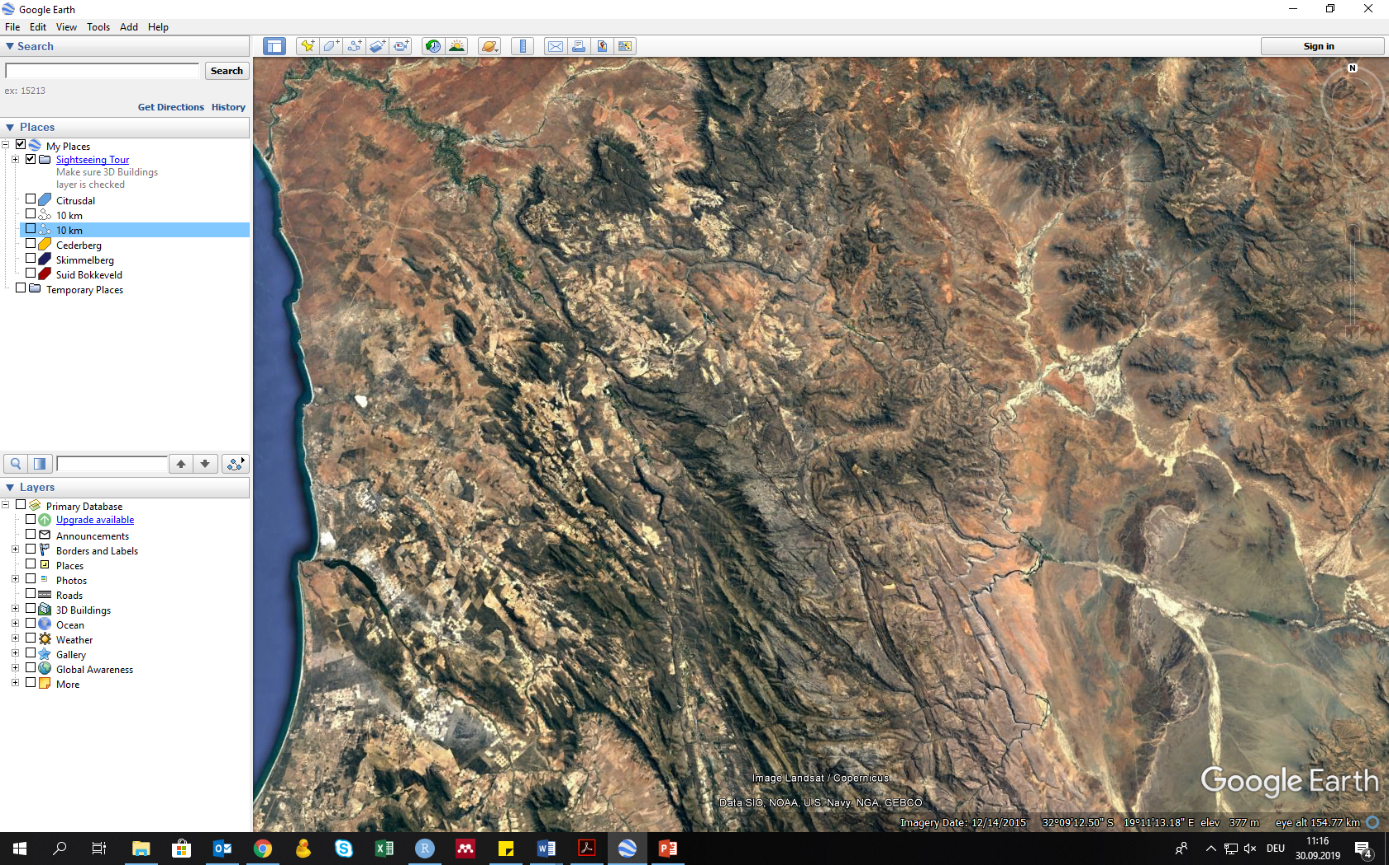

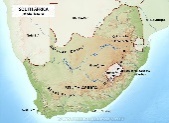

**Suid Bokkeveld**

**Cederberg**

**Skimmelberg**

**Citrusdal**

**10 km**

**Figure S2 Calinski-Harabasz (CH) scores of the predicted clusters of root nodule communities of rooibos based on Jensen-Shannon distances.** The highest CH score indicates the most optimal number of k-clusters in which a data distribution can be classified.

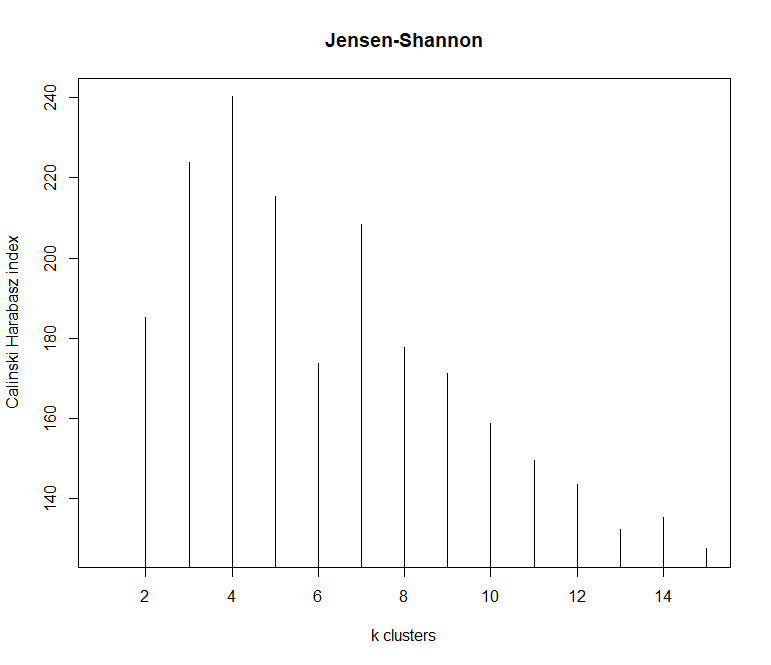

k-clusters

2

4

6

8

10

12

14

Calinski-Harabasz index

140

160

180

200

220

240

**Figure S3 Principal components analysis (PCA) of the stoichiometric leaf trait variation of cultivated and wild rooibos in this study.** Arrow overlays depict the direction of the variation explained by the leaf stoichiometric traits that varied across predicted root nodule communities of rooibos. A 650bp fragment of the *psbA-trnH* chloroplastic gene marker was used to characterize rooibos genotypes. Genotypes A and G were the only ones represented in more than 10 plants in this study and are thus the only ones shown.

PC2 (32.35%)

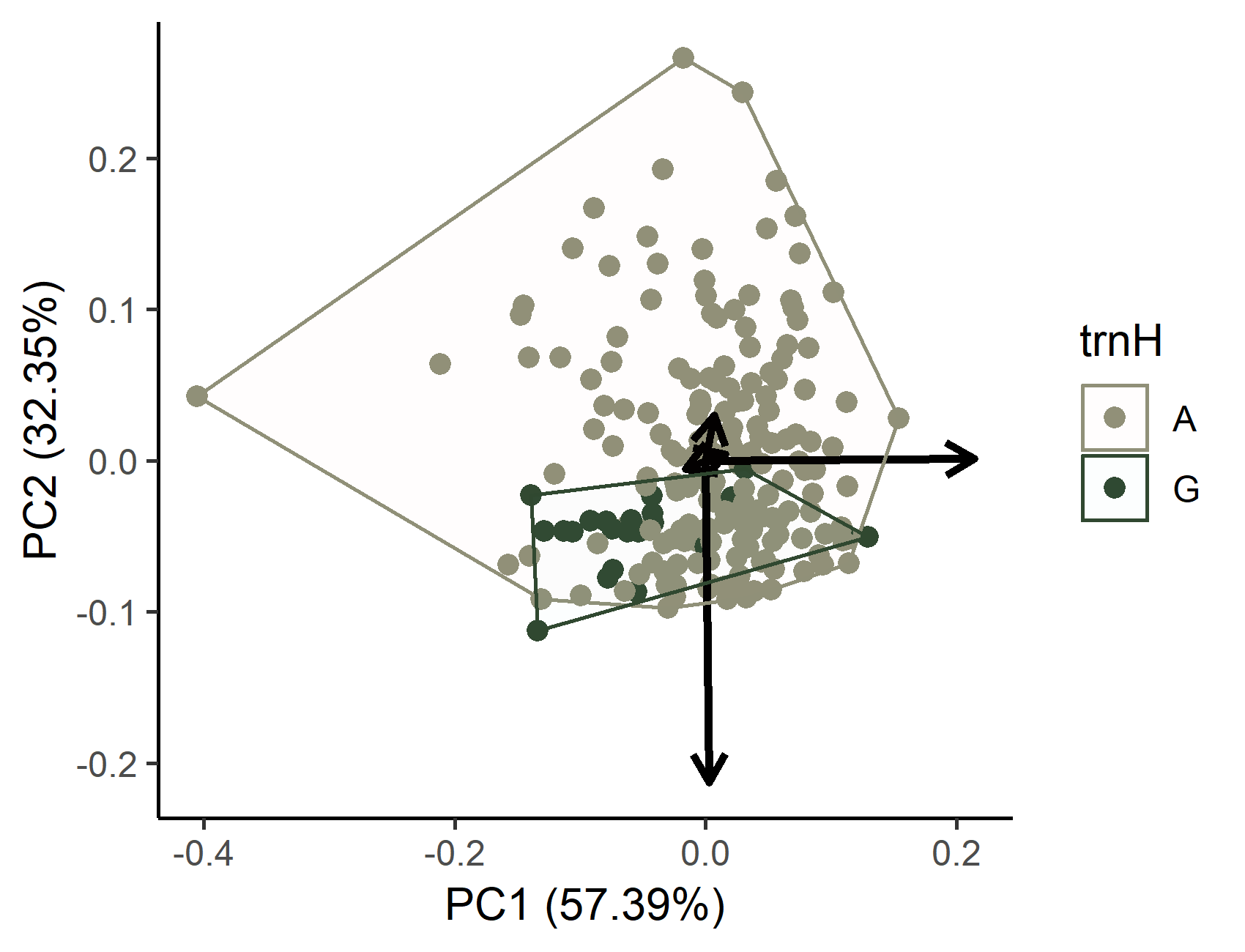

PC1 (57.39%)

0.2

-0.2

-0.4

0

0

0.1

0.2

-0.2

-0.1

^13^C (‰)

^15^N (‰)

K (mg g^-1^)

C:N ratio

N (mg g^-1^)

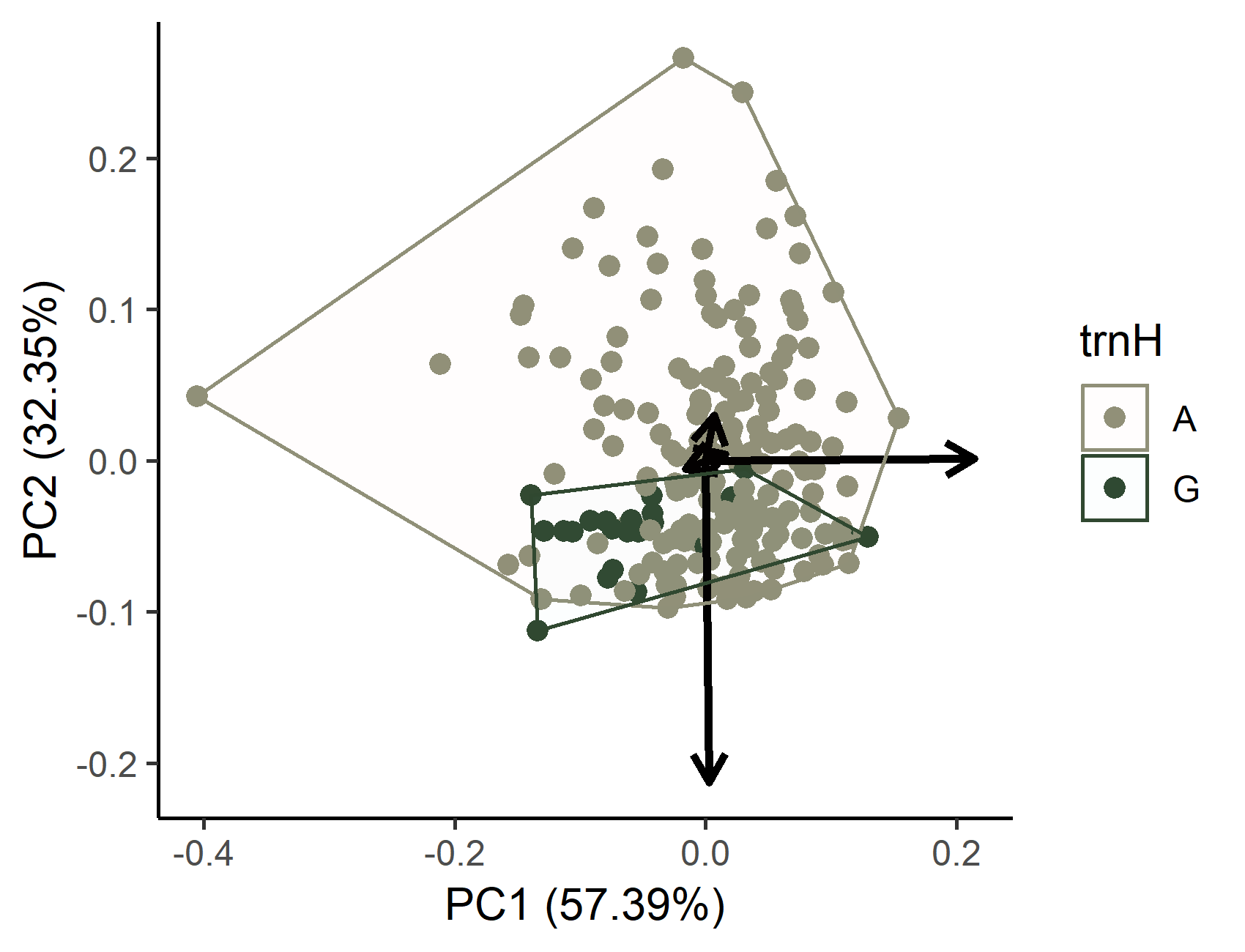

A

G

**Rooibos genotype (*psbA-trnH* based)**

**Figure S4 Linear regressions showing no relationship between soil N and K concentrations and N:P ratios and root nodule community dissimilarity based on *gyrB* sequence differences.** Dots depict distances between centroids of Bray-Curtis dissimilarity values between root nodule communities.

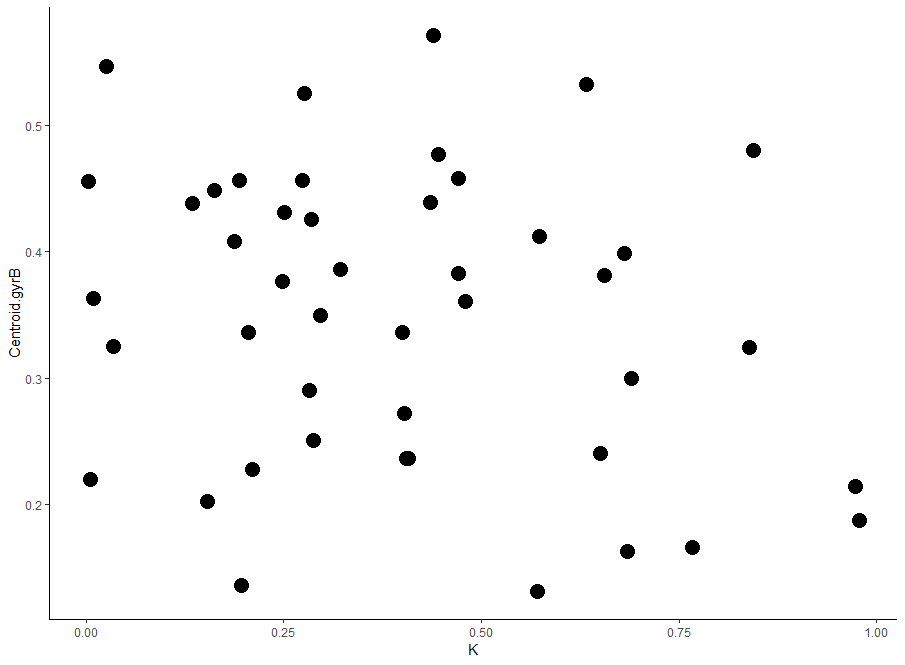

Soil K (g kg^-1^)

0

0.25

0.50

0.75

1.00

0.2

0.3

0.4

0.5

Distance between centroids of root nbodule communities

ns

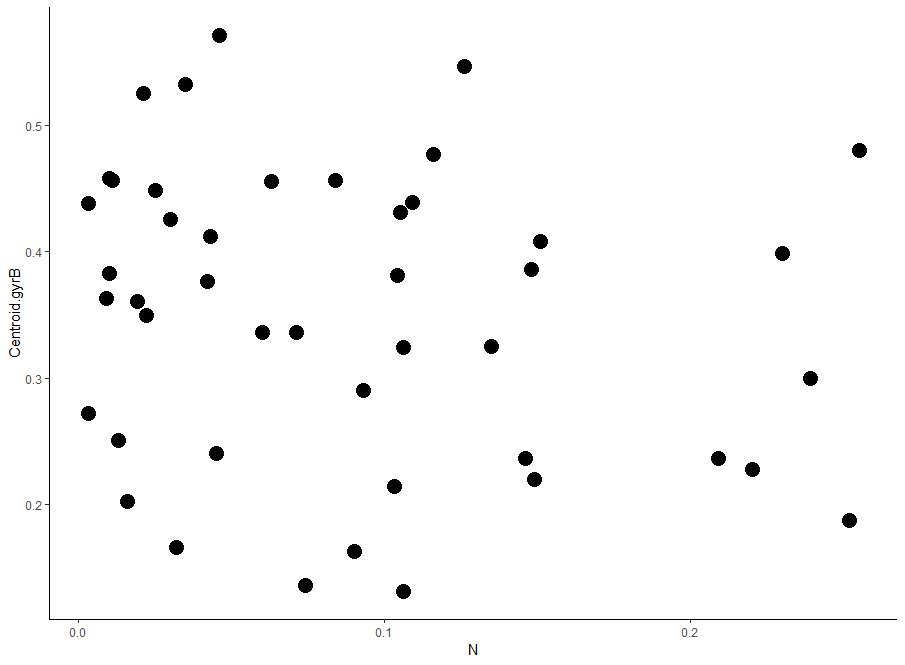

Soil N (g kg^-1^)

0.1

0.2

0

0.2

0.3

0.4

0.5

Distance between centroids of root nodule communities

ns

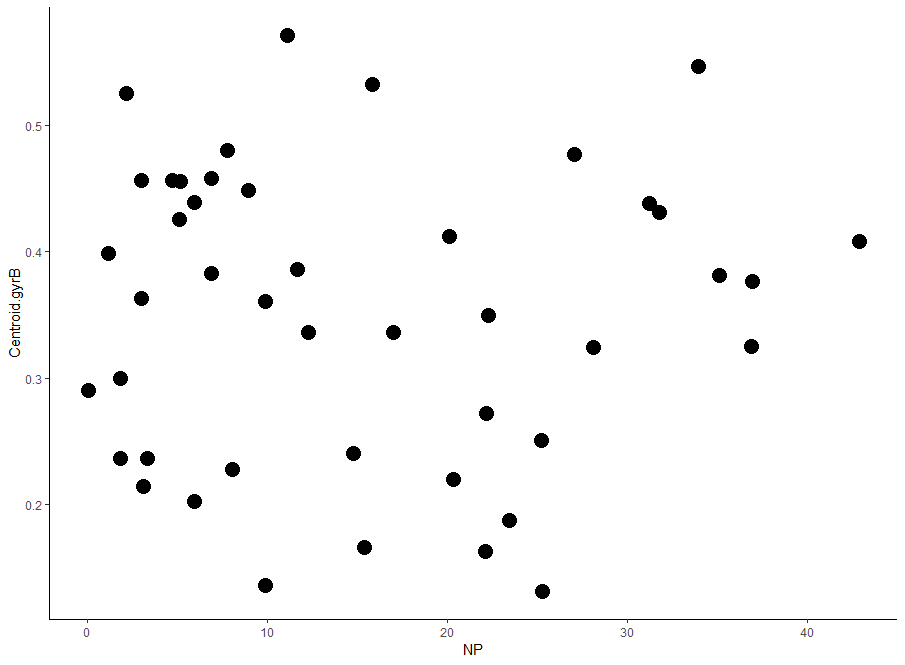

0.2

0.3

0.4

0.5

Soil N:P ratio

0

10

20

30

40

ns

**Figure S5 Linear regressions showing no relationship between geographical distance and rooibos root nodule community dissimilarity based on *gyrB* sequence differences within the Suid Bokkeveld (South Africa).** (A) Absence of a distance-decay relationship of community similarity from root nodule communities obtained from wild rooibos populations. (B) Absence of a distance-decay relationship of community similarity between root nodule communities obtained from cultivated rooibos populations. Dots depict distances between centroids of Bray-Curtis dissimilarity values between root nodule communities.

**A**

**B**

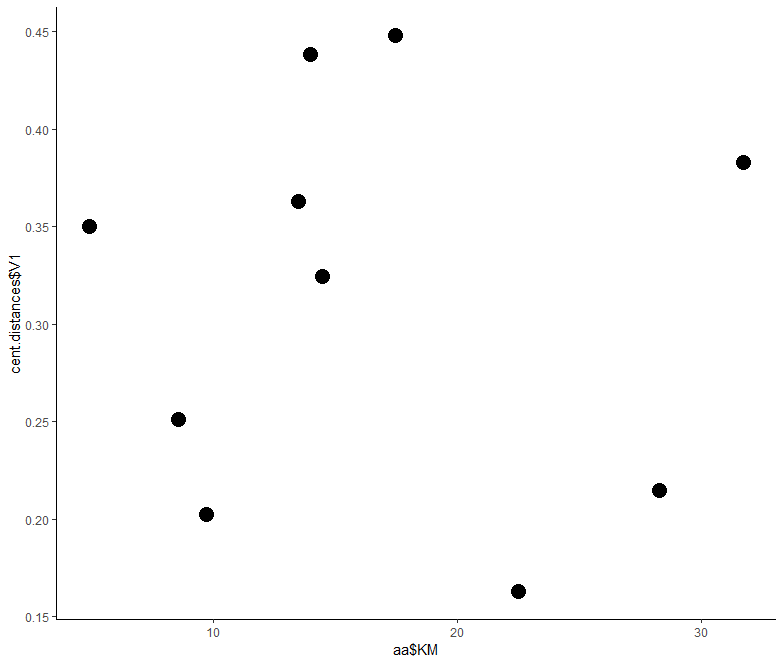

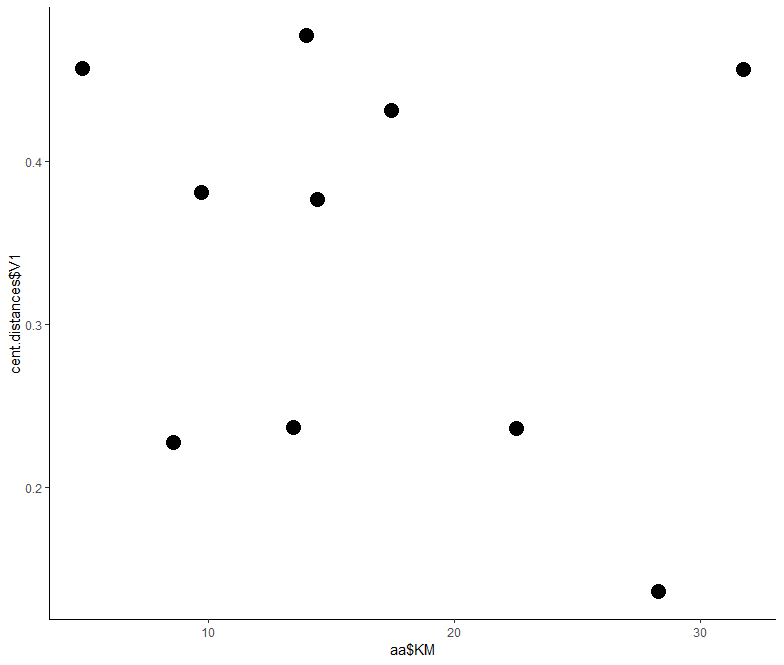

0.2

0.3

0.4

10

20

30

Geographical distance between locations (km)

ns

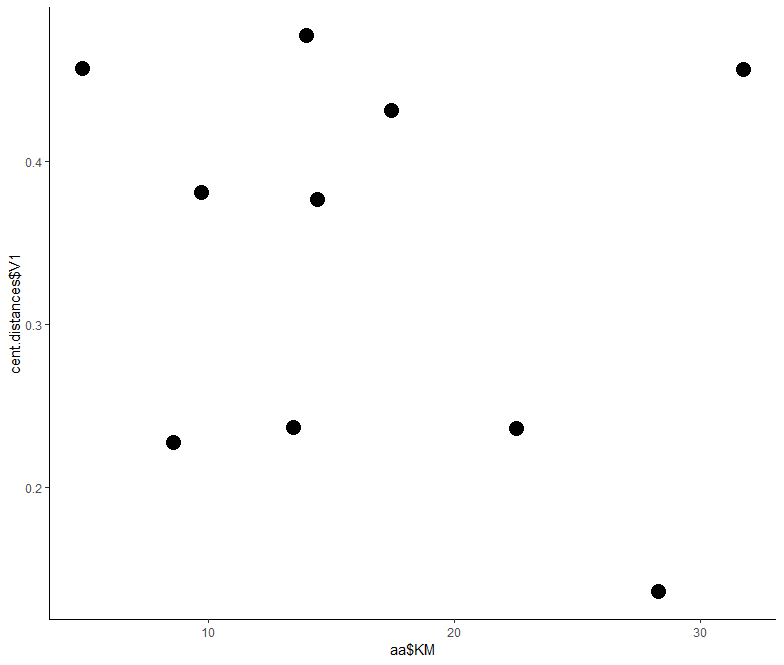

0.2

0.3

0.4

10

20

30

Distance between centroids of root nodule communities

Geographical distance between locations (km)

ns

**Figure S6 Linear regressions showing no relationship between geographical distance and rooibos root nodule community dissimilarity based on *nodA* functional gene sequence differences in the rooibos farming range.** (A) Absence of a distance-decay relationship of community similarity from root nodule communities obtained from wild rooibos populations. (B) Absence of a distance-decay relationship of community similarity between root nodule communities obtained from cultivated rooibos populations. Dots depict distances between centroids of Bray-Curtis dissimilarity values between root nodule communities.

**B**

**A**

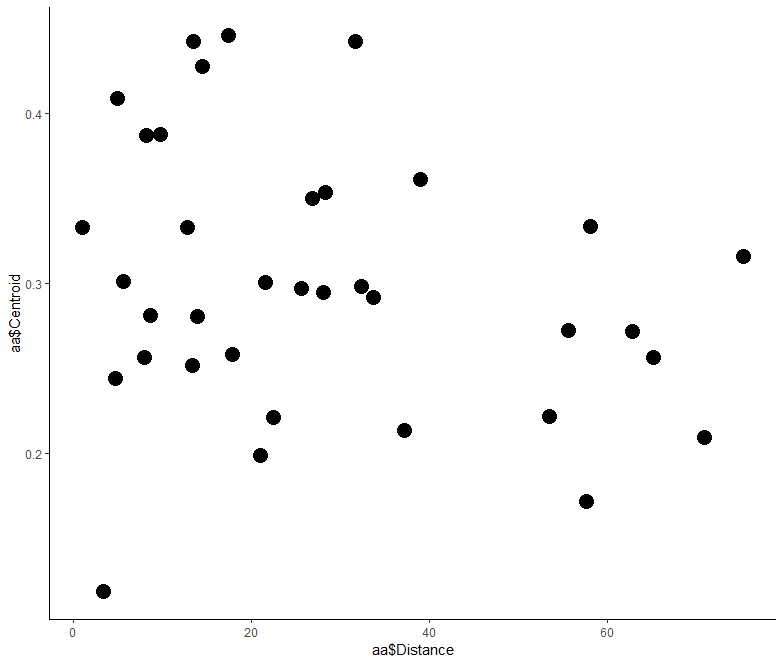

0.2

0.3

0.4

20

Geographical distance between locations (km)

40

60

0

ns

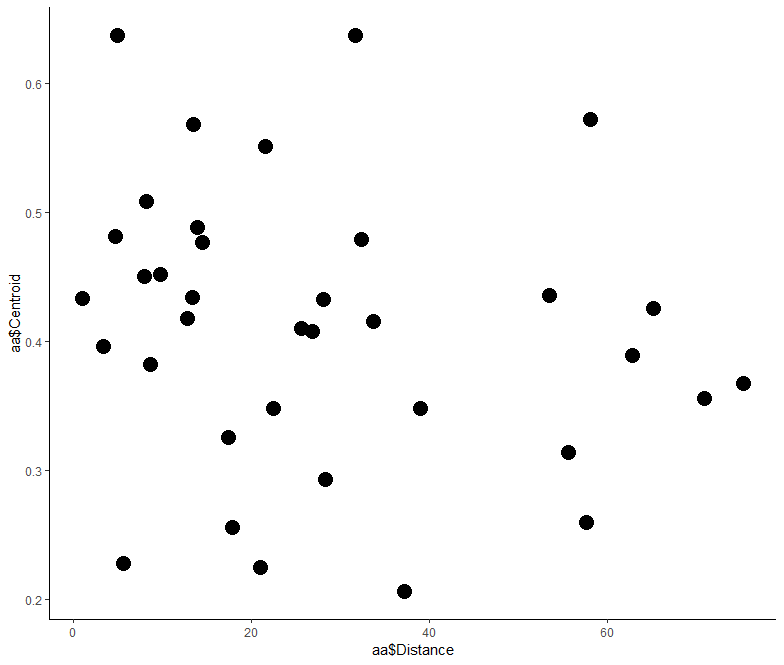

20

Geographical distance between locations (km)

40

60

0

0.2

0.4

0.6

Distance between centroids of root nodule communities

0.3

0.5

ns

**Figure S7 Map of the Suid Bokkeveld (South Africa) with the proportion of the four root nodule community classes predicted in each of the six organic rooibos farms sampled.** The top panel depicts the proportion of samples from the six different farms that belongs to each of the four predicted root nodule community classes. Root nodule communities were described using a fragment of the *gyrB* taxonomic gene marker. Farms: BF, Blomfontein; DB, Dobbelarskop; LK, Landsklof; MA, Matarakoppies; SO, Sonderwaterkraal.

MA

BF

DB

LK

MK

SO

**Farm**

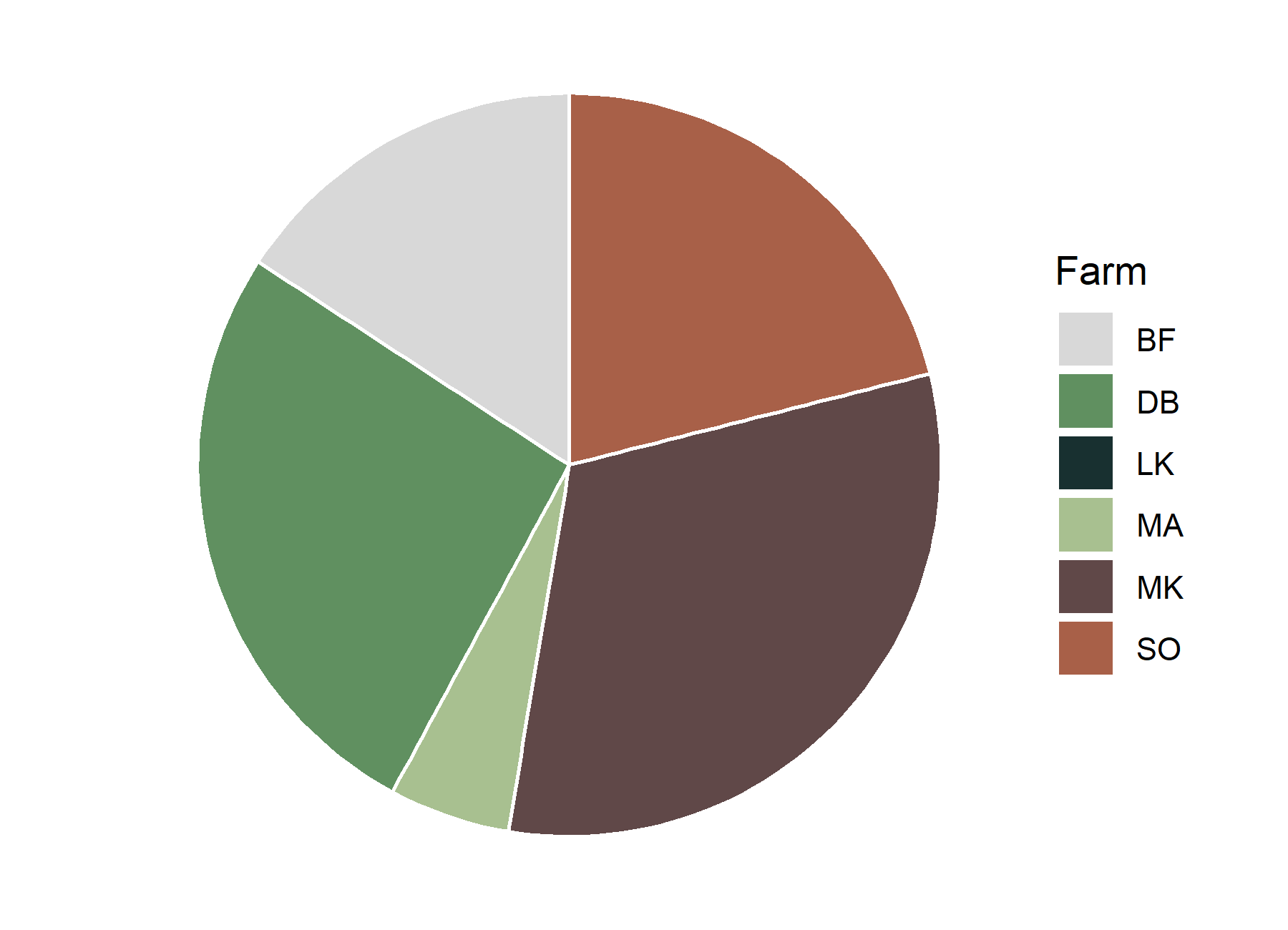

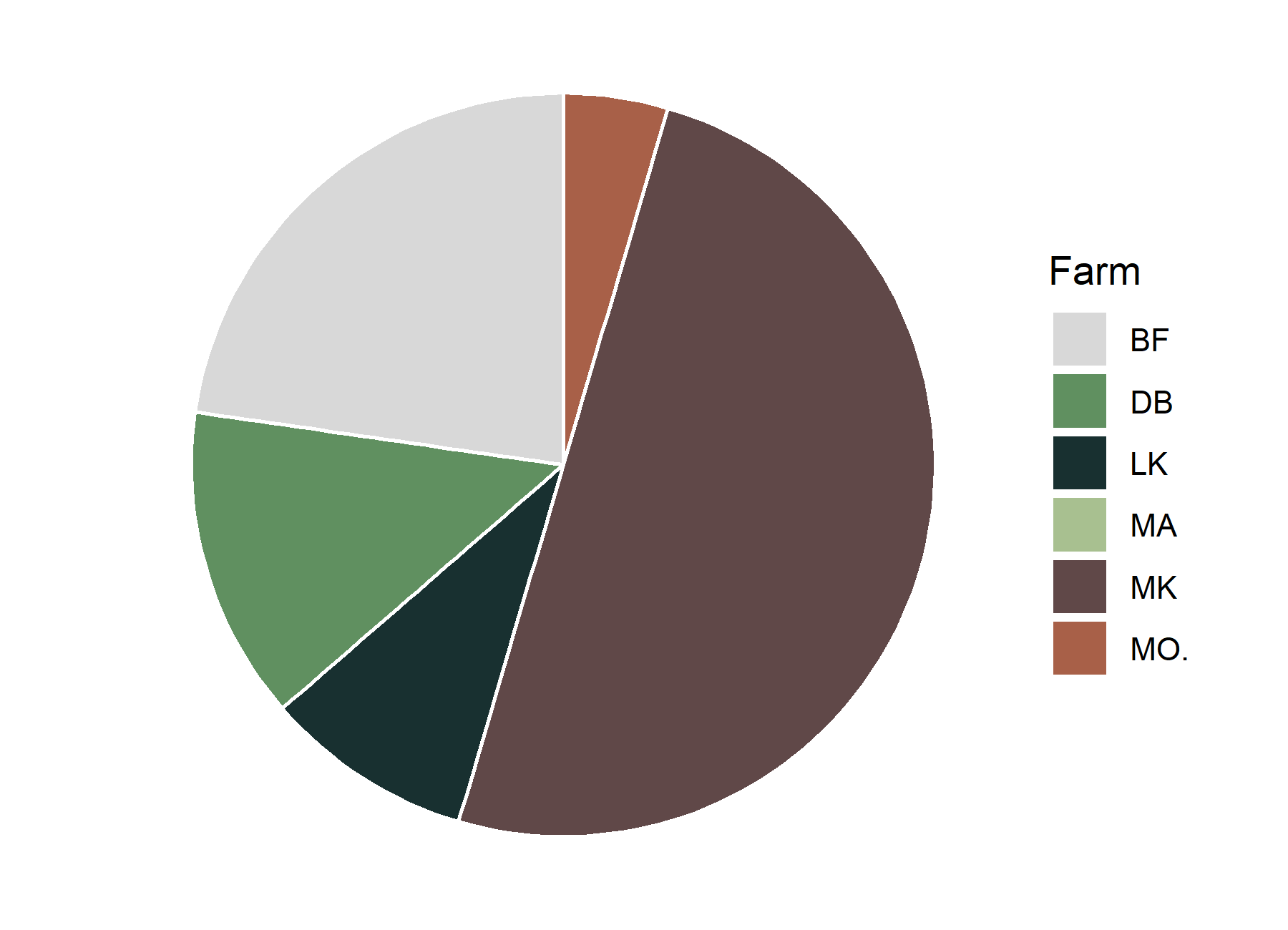

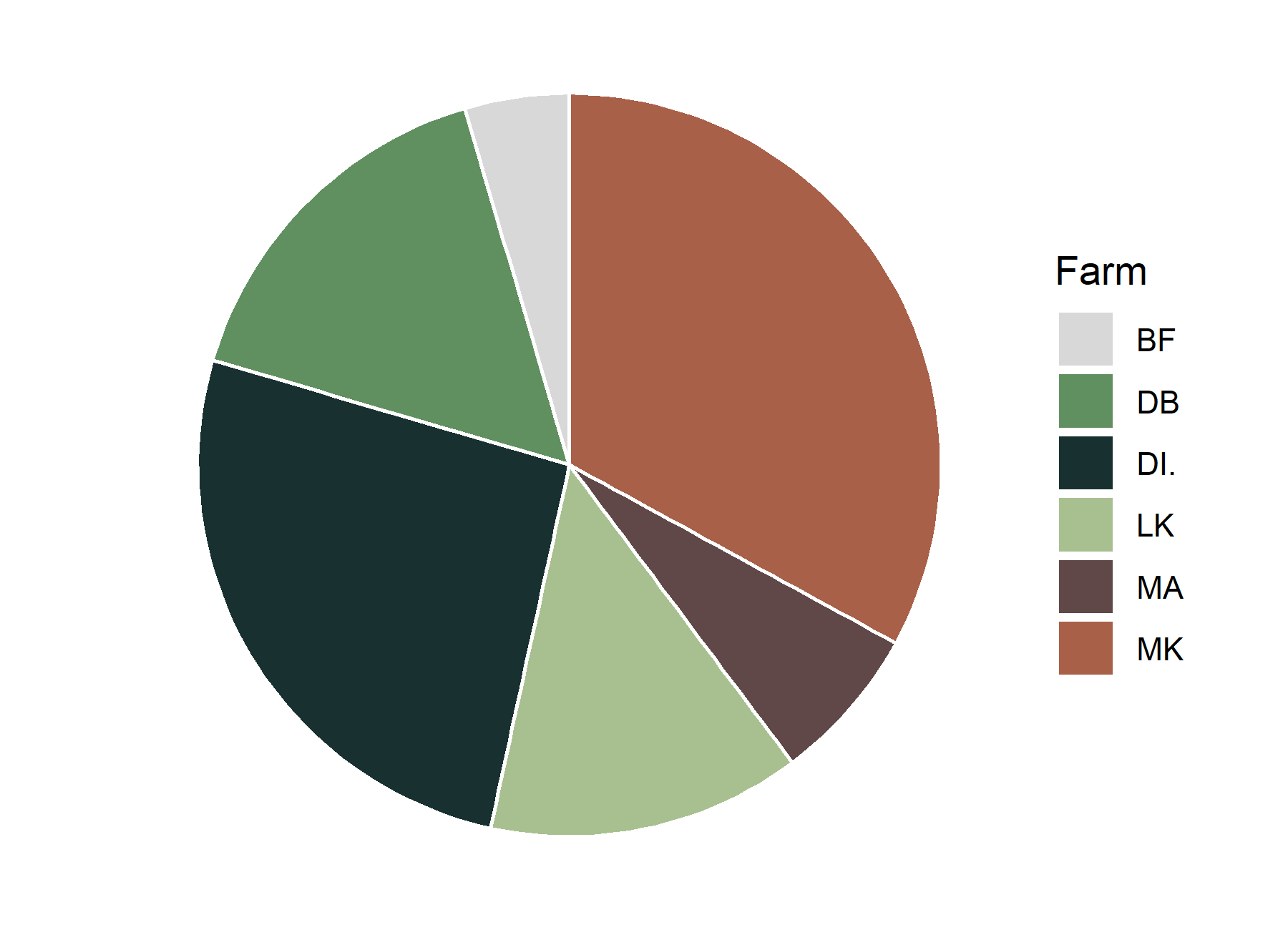

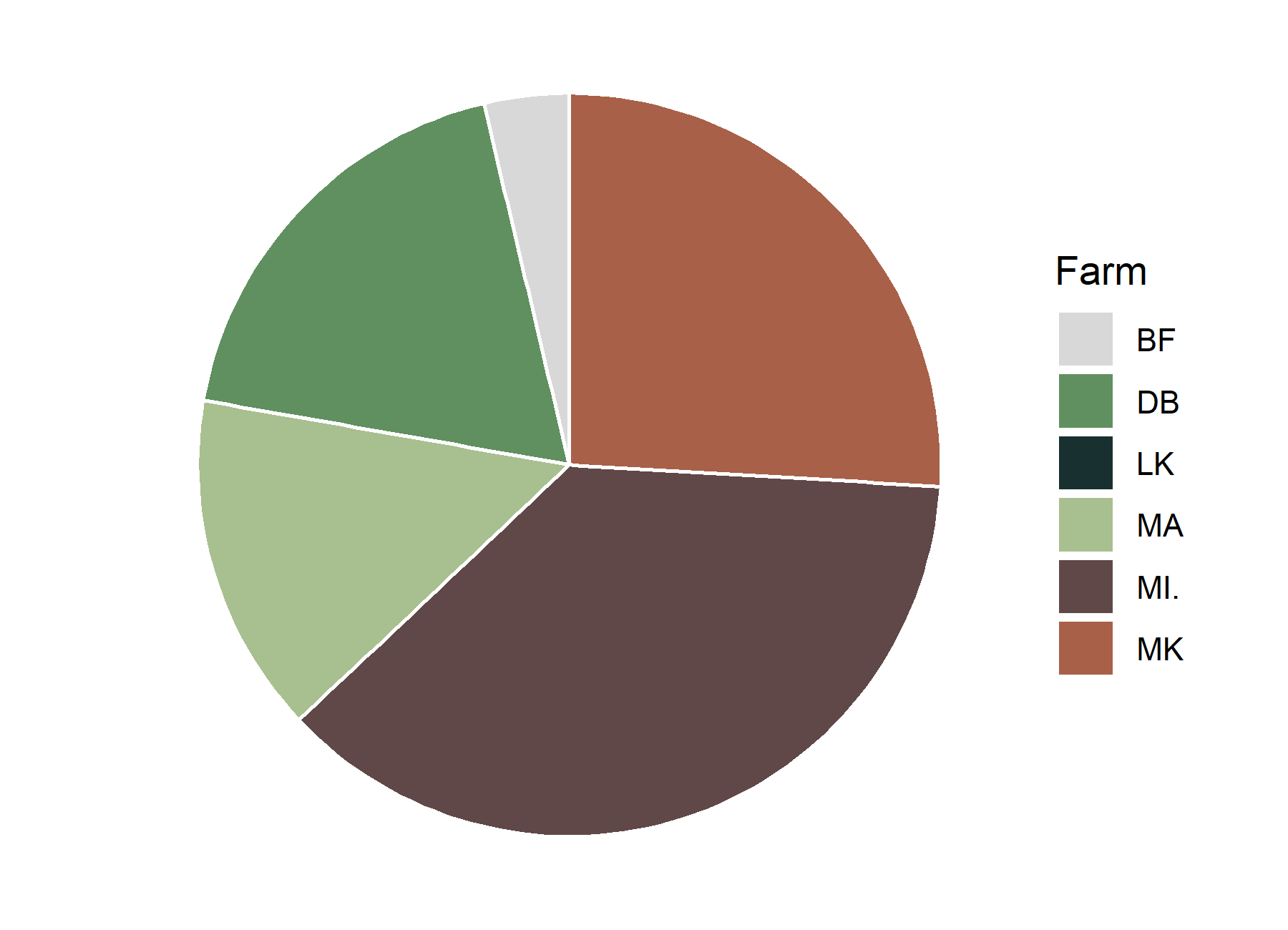

1

2

3

4

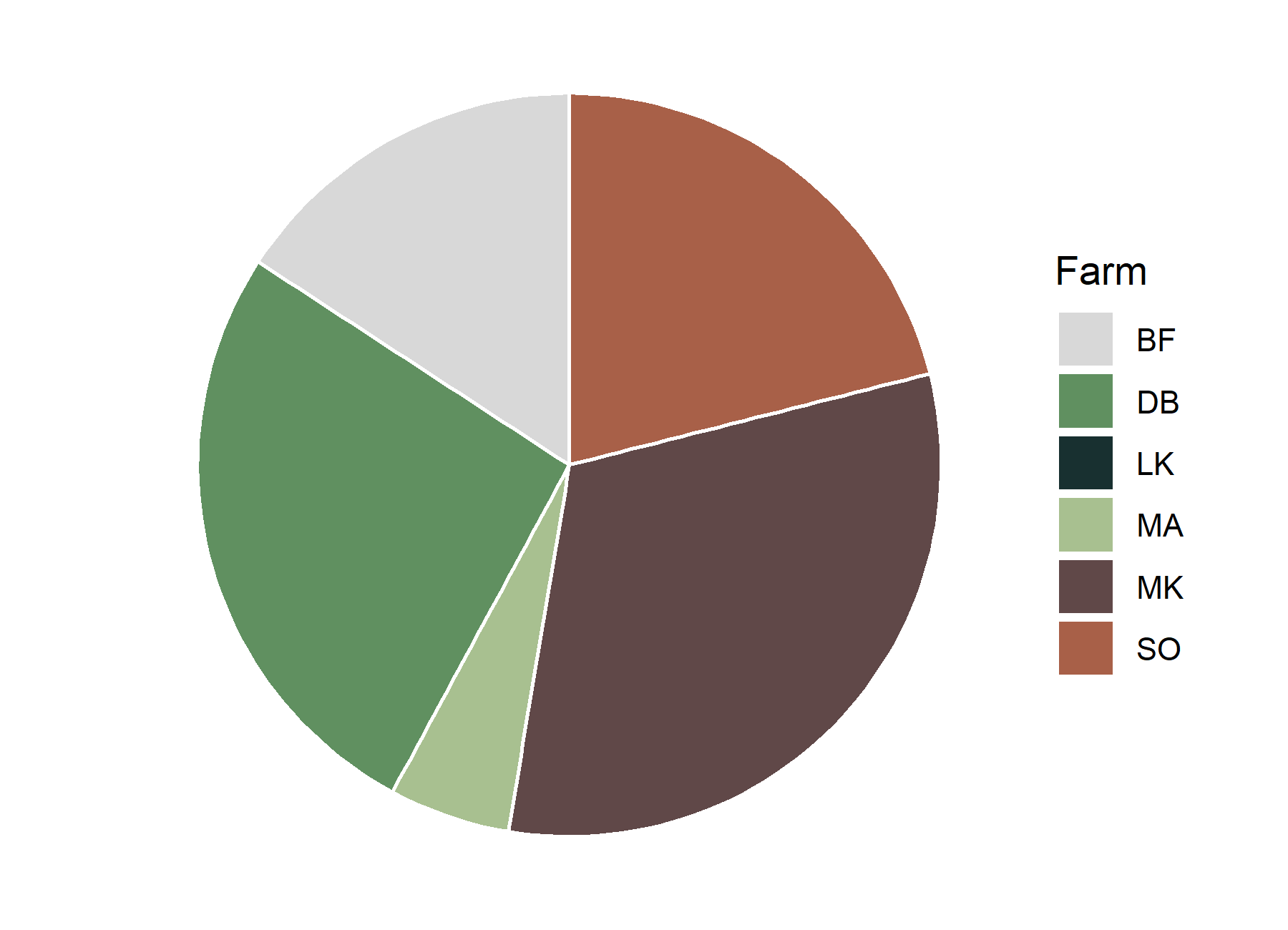

**Community class**

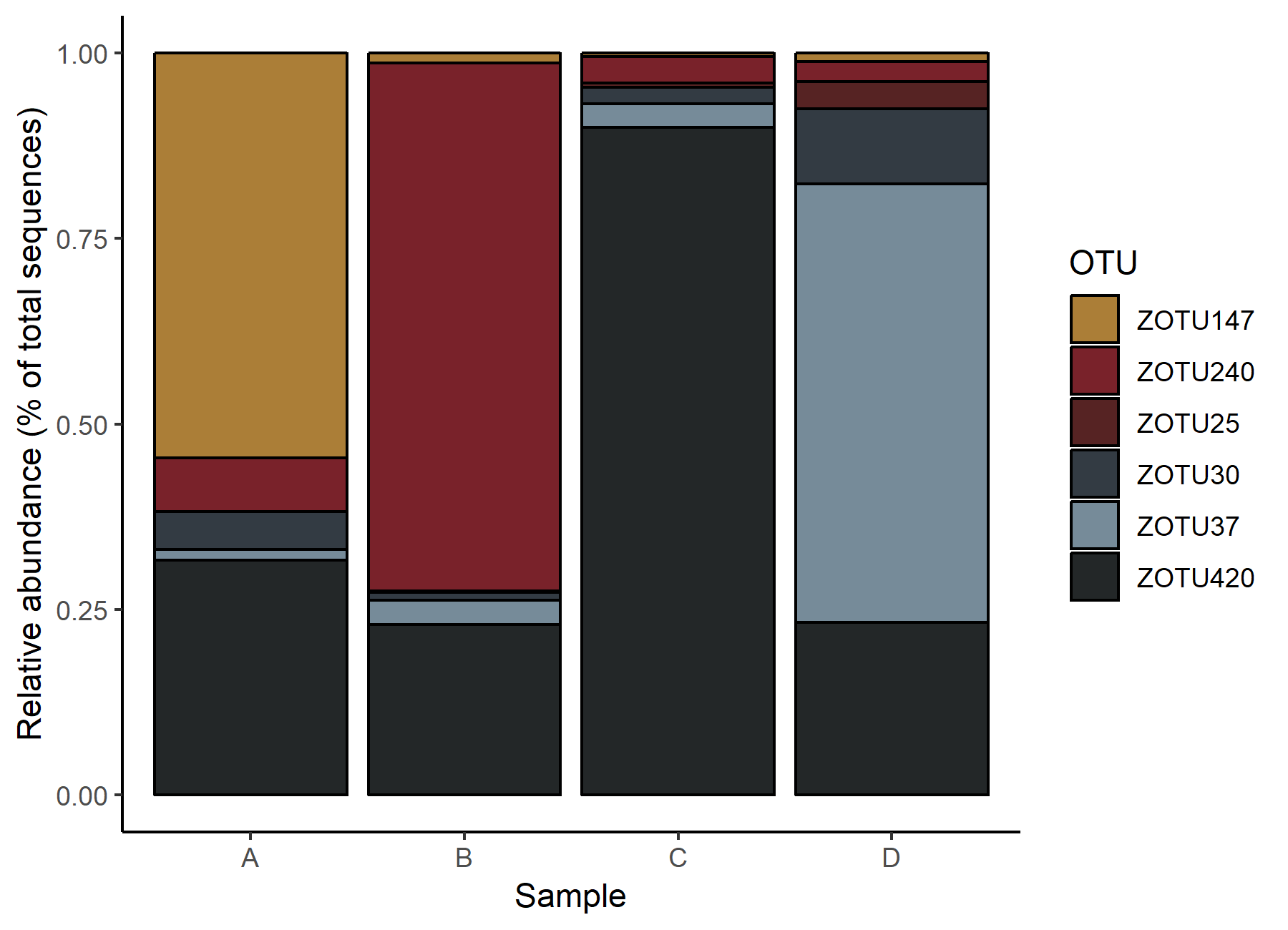

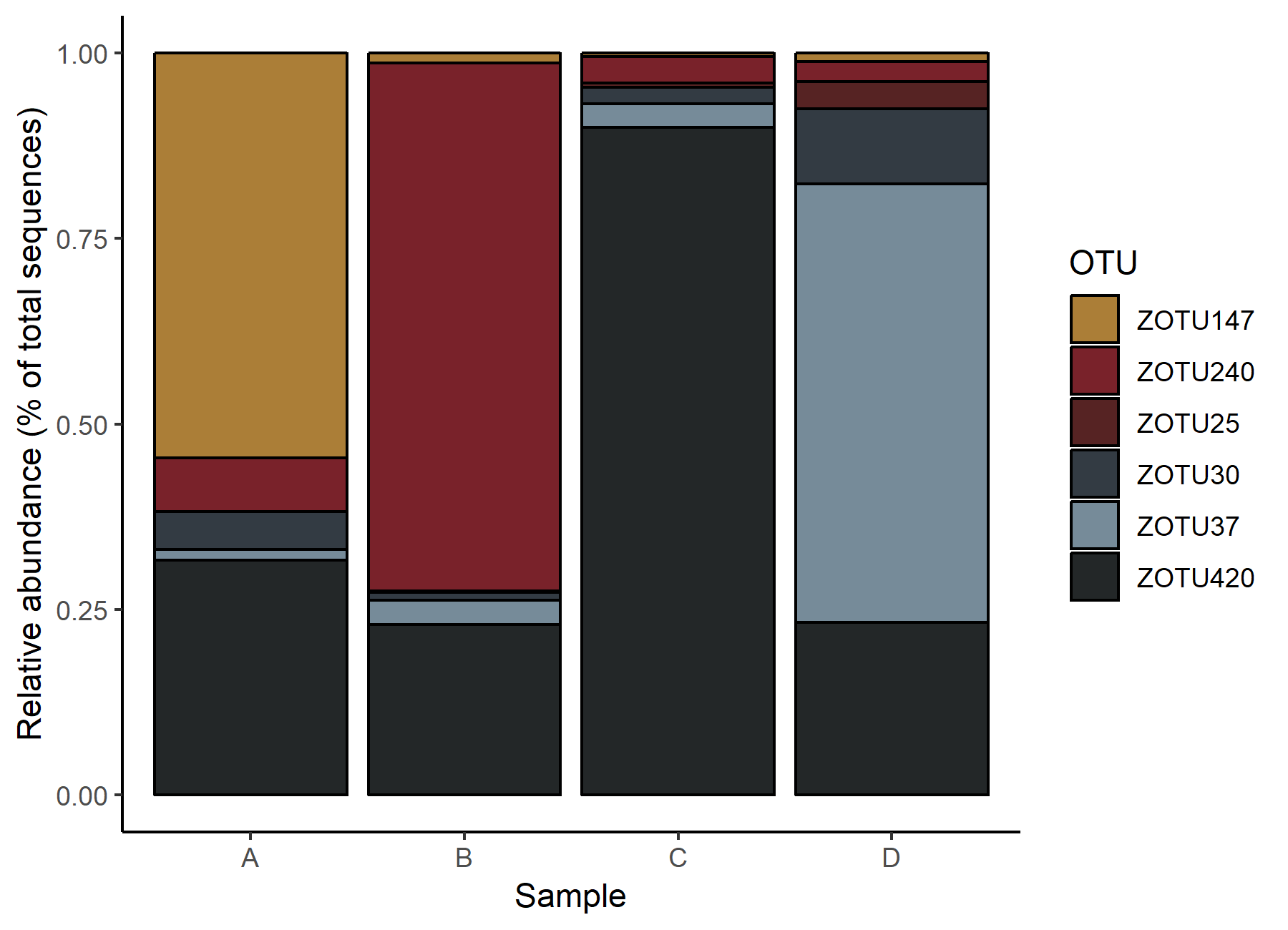

1

2

3

4

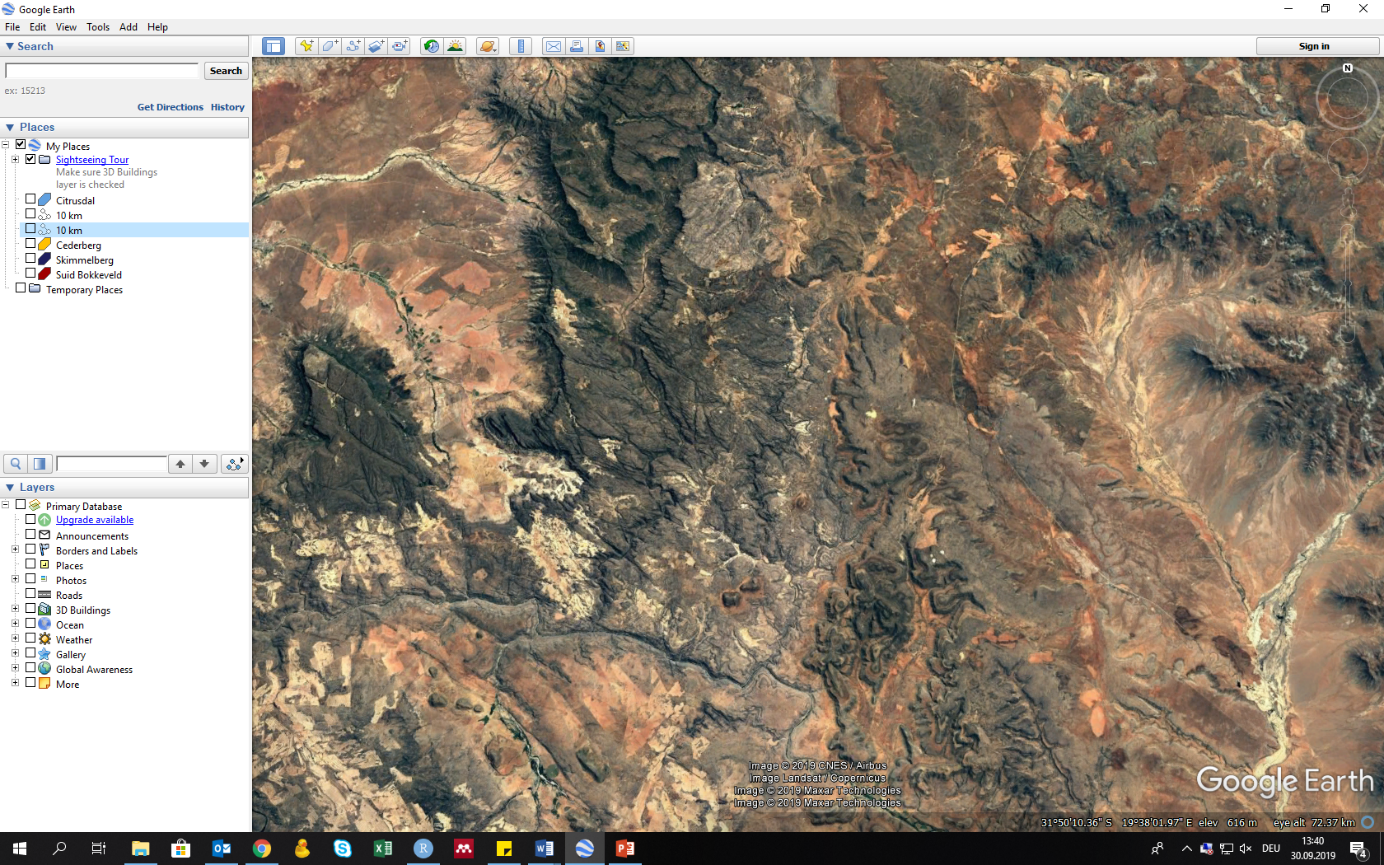

**5 km**

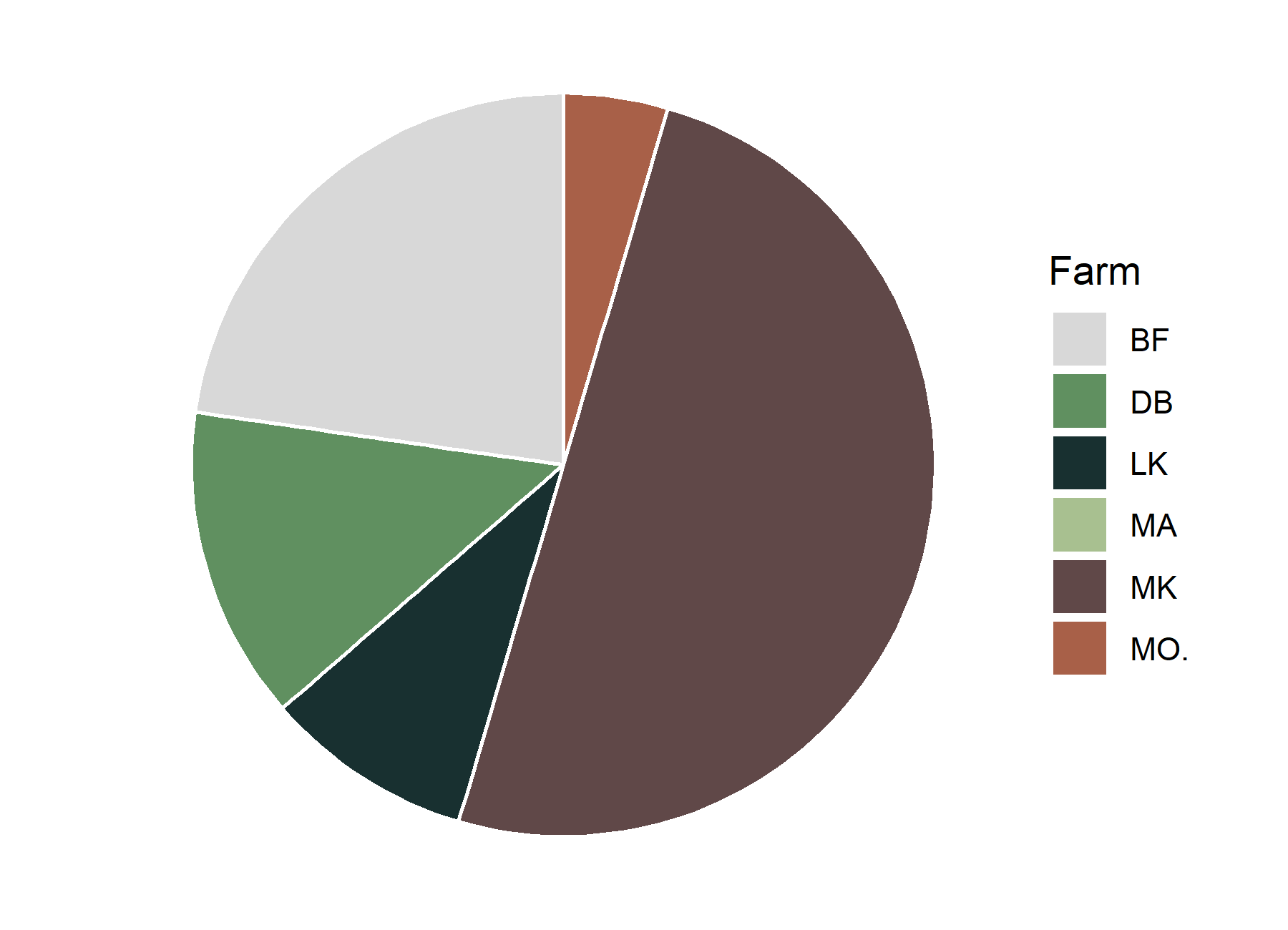

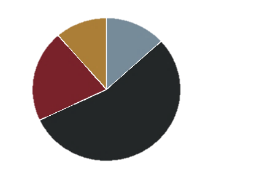

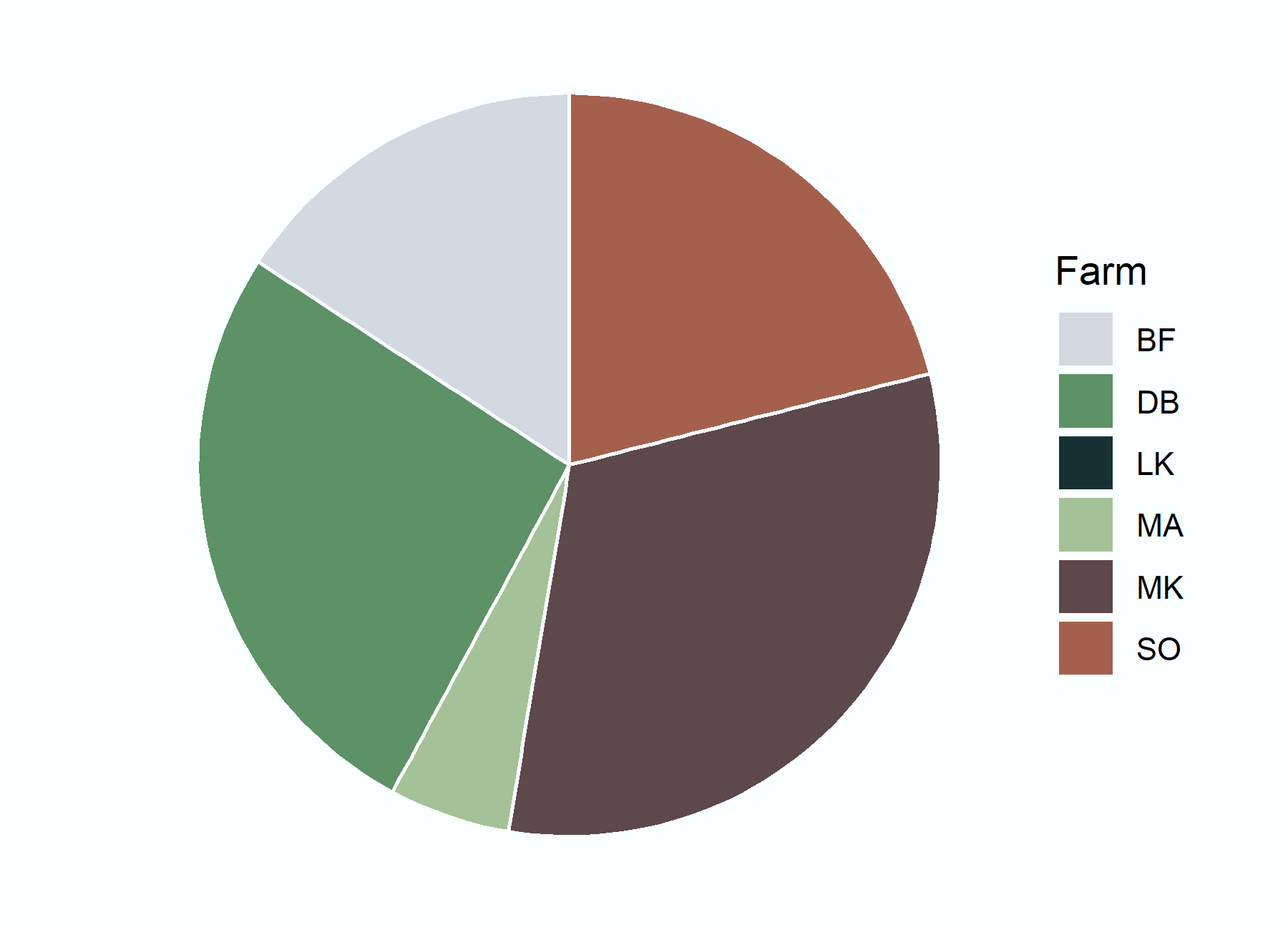

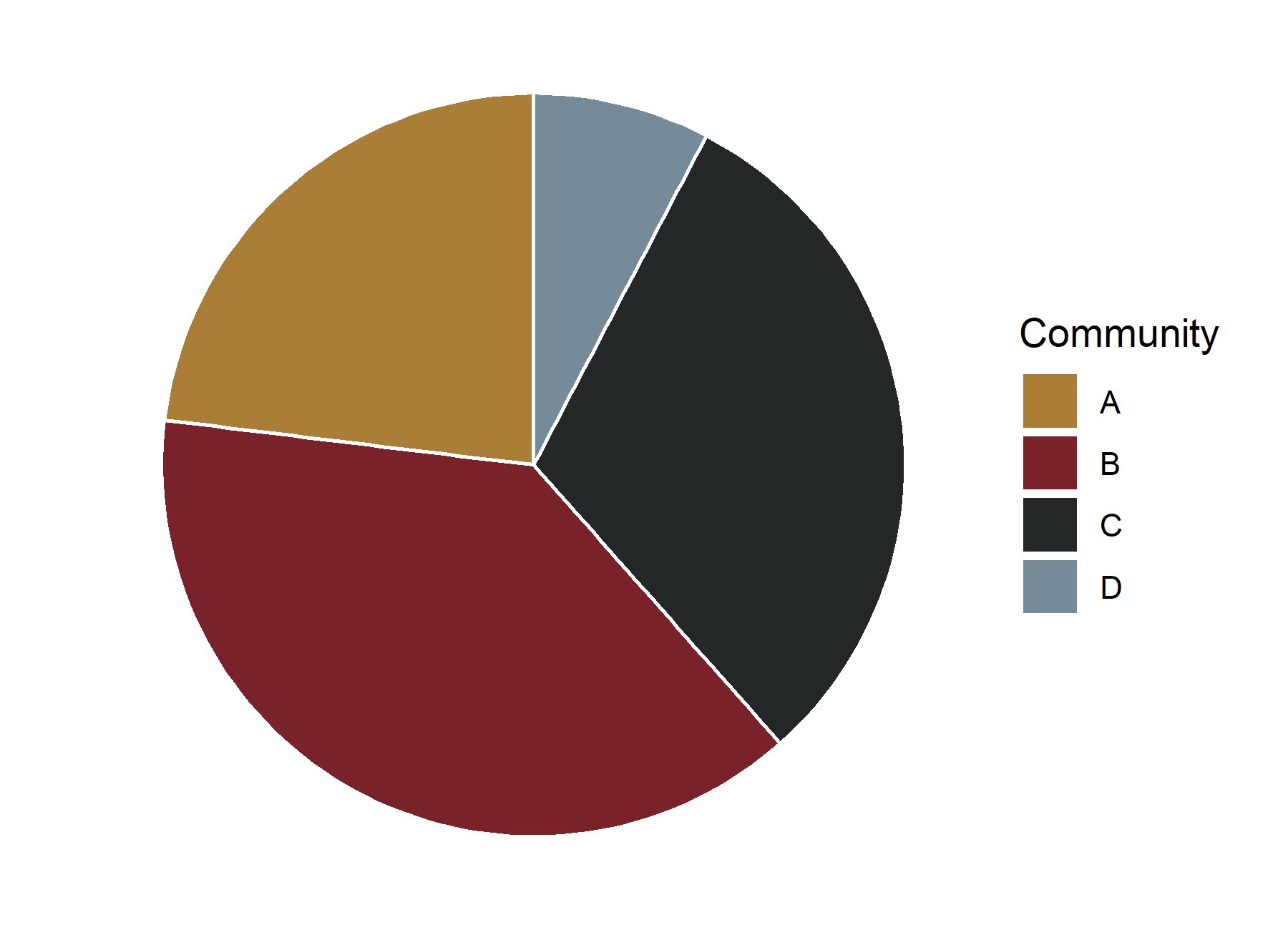

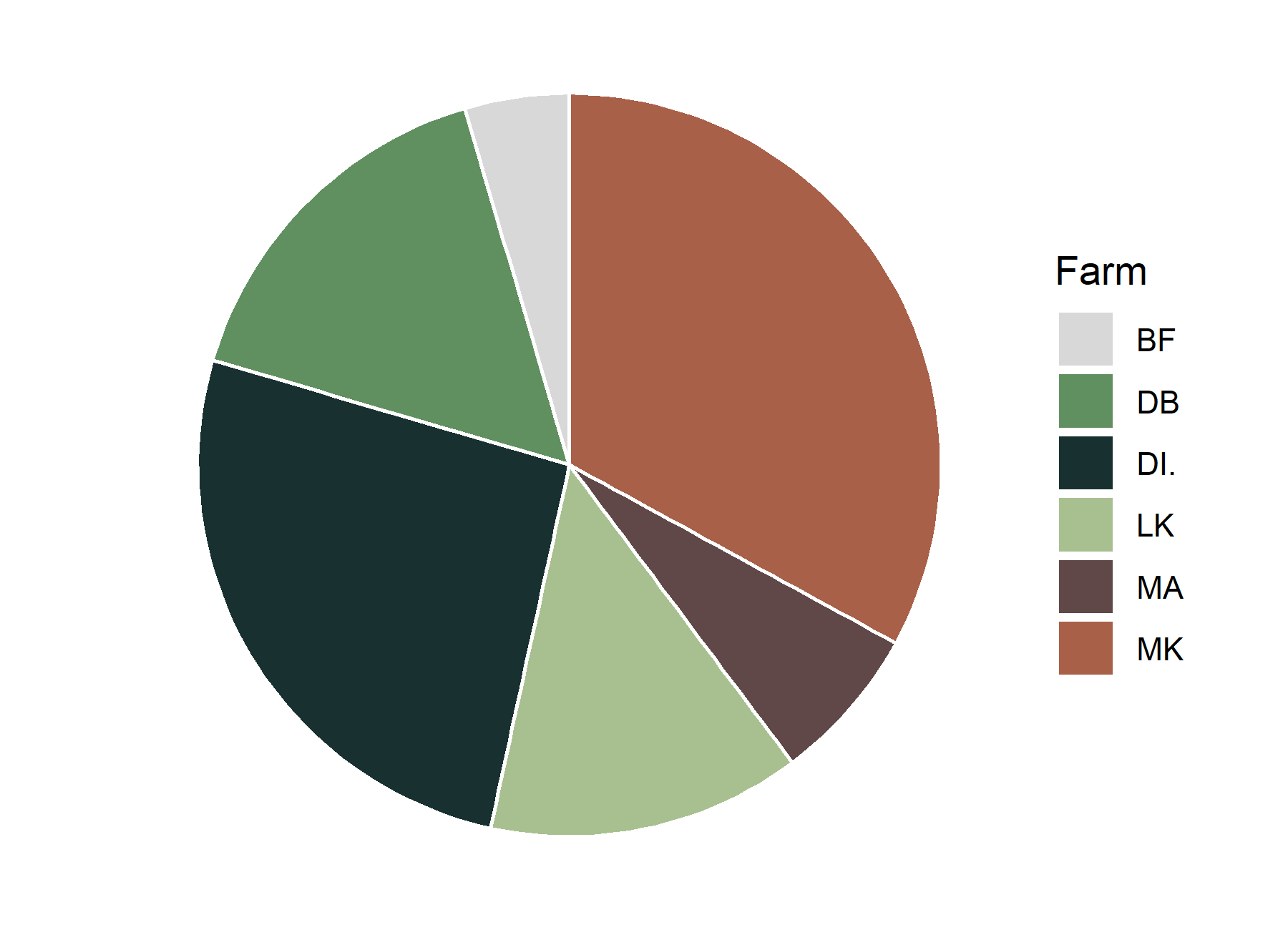

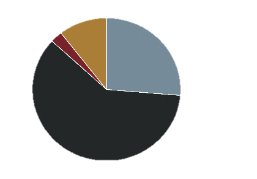

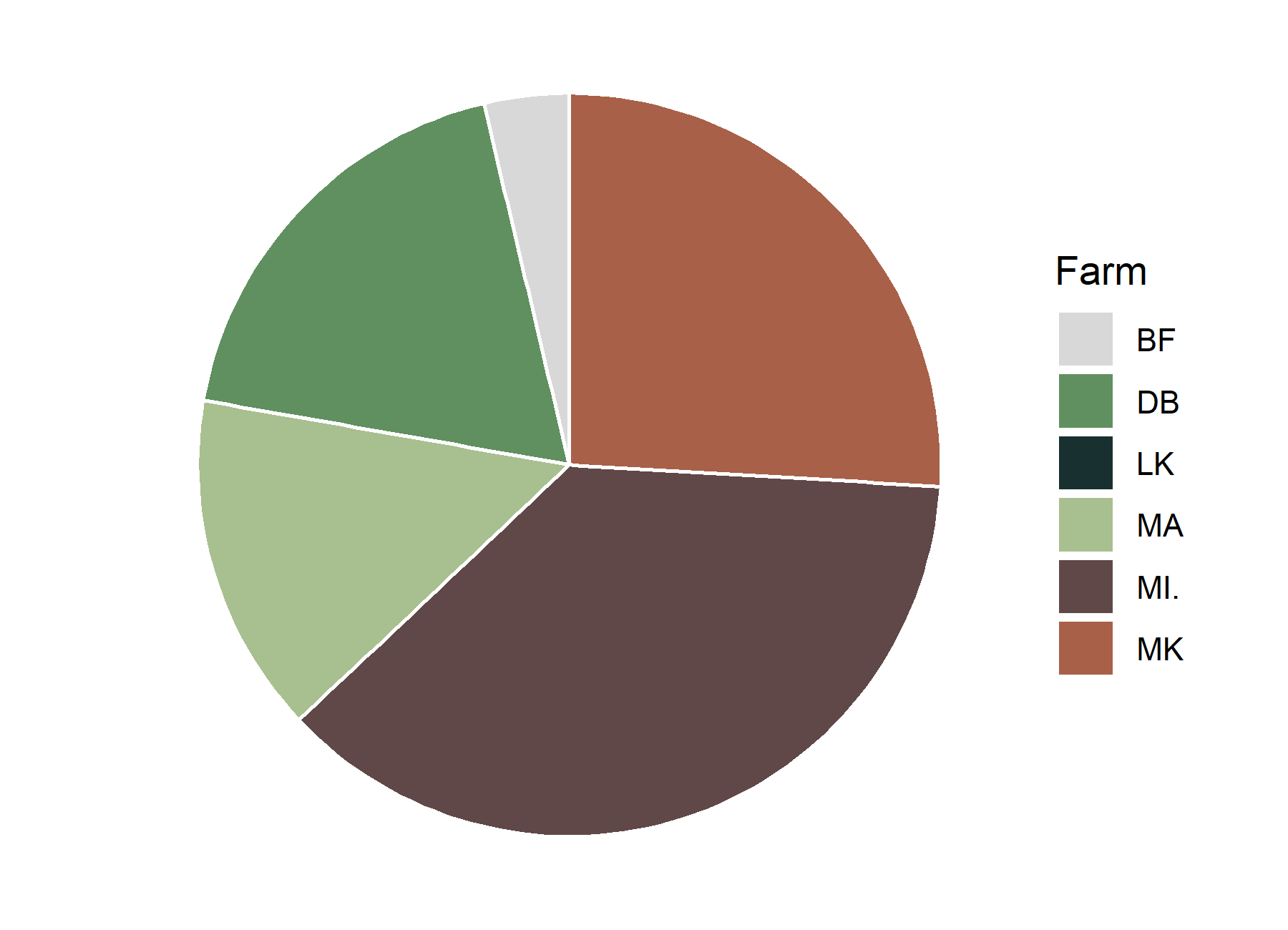

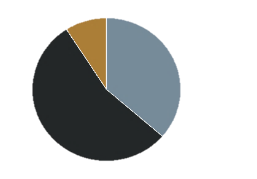

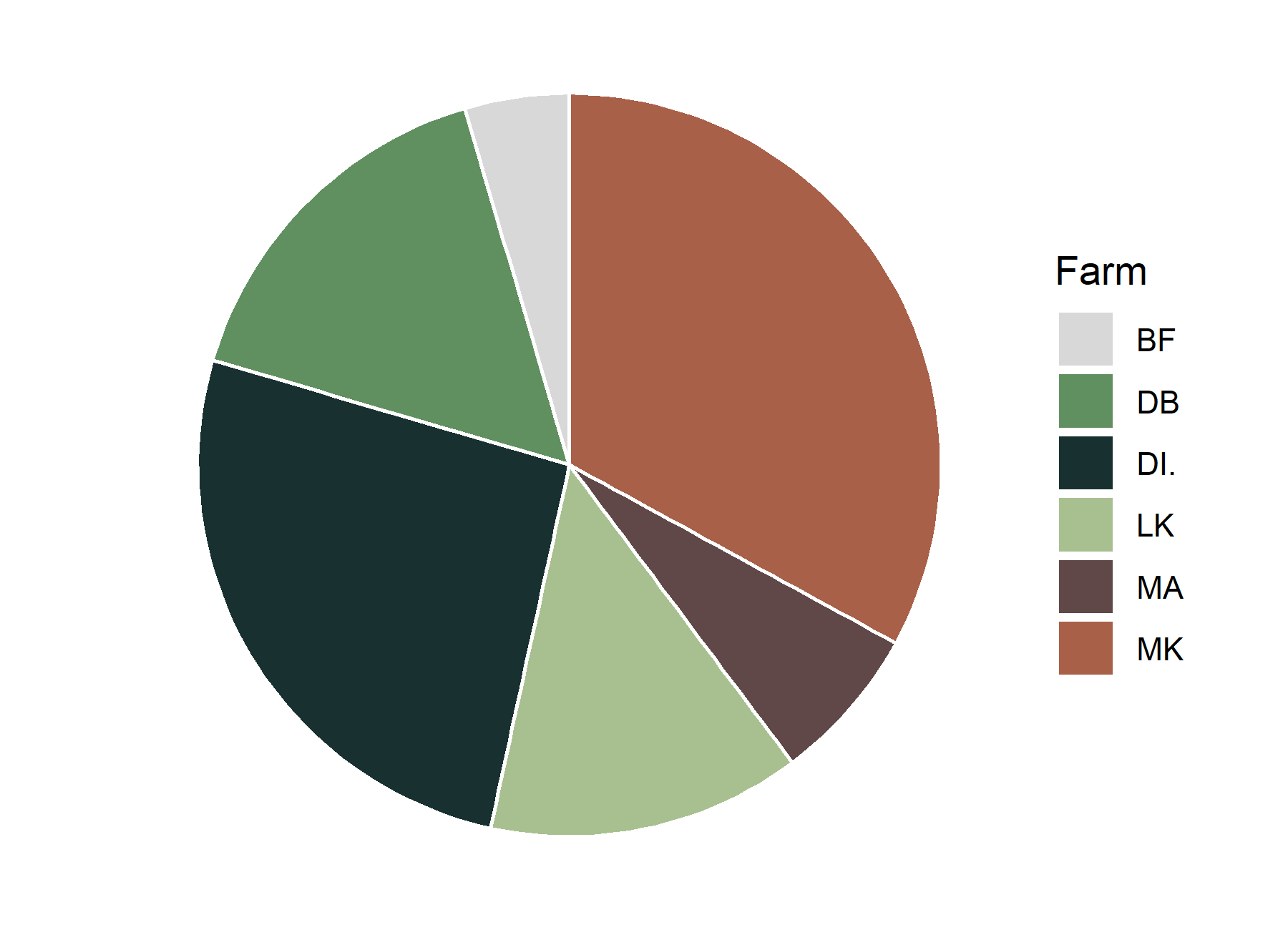

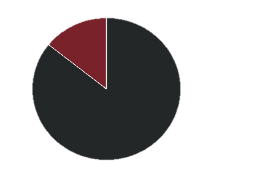

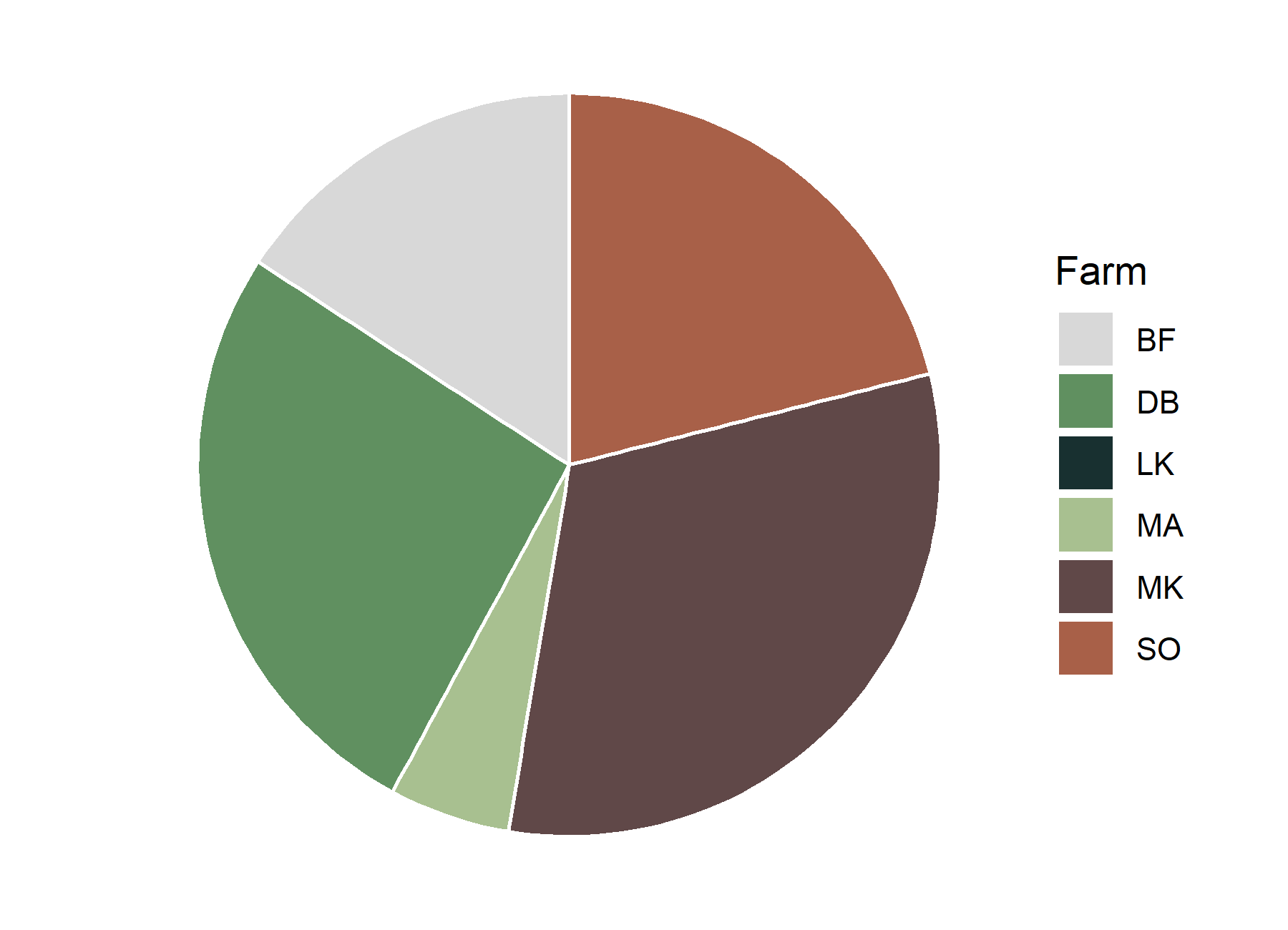

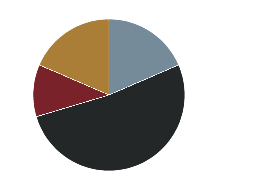

**Table S1 Nutrient concentrations and pH of soils collected from around rooibos plants in the Suid Bokkeveld (South Africa).** Locations/farms: Blo, Blomfontein; Dob, Dobbelarskop; Lan, Landsklof; Mat, Matarakoppies; Mel, Melkkraal.

| *Soil properties* | *Cultivated soil (plantations)*  *(Farms)* | | | | | *Uncultivated soil (wild populations)*  *(Farms)* | | | | |
| --- | --- | --- | --- | --- | --- | --- | --- | --- | --- | --- |
|  | *Blo* | *Dob* | *Lan* | *Mat* | *Mel* | *Blo* | *Dob* | *Lan* | *Mat* | *Mel* |
| pH (CaCl_2_) | 5.78 | 5.76 | 6.15 | 5.53 | 5.55 | 5.89 | 5.48 | 5.51 | 5.60 | 5.29 |
| N (g kg^-1^) | 0.11 | 0.13 | 0.11 | 0.14 | 0.22 | 0.26 | 0.37 | 0.15 | 0.16 | 0.22 |
| P (g kg^-1^) | 0.020 | 0.001 | 0.003 | 0.010 | 0.007 | 0.005 | 0.030 | 0.007 | 0.010 | 0.020 |
| K (g kg^-1^) | 0.38 | 0.53 | 0.24 | 0.54 | 1.22 | 0.57 | 1.22 | 1.01 | 0.82 | 0.81 |
| δ^15^N (‰) | 20.40 | 19.75 | 23.68 | 22.29 | 14.52 | 10.87 | 10.31 | 20.26 | 19.31 | 13.91 |
| N:P ratio | 4.98 | 10.94 | 36.17 | 13.91 | 33.06 | 47.84 | 12.74 | 20.80 | 16.07 | 10.91 |
| Ca (g kg^-1^) | 0.06 | 0.11 | 0.06 | 0.05 | 0.12 | 0.14 | 0.16 | 0.06 | 0.06 | 0.08 |
| Mg (g kg^-1^) | 0.10 | 0.17 | 0.12 | 0.07 | 0.29 | 0.17 | 0.35 | 0.12 | 0.07 | 0.15 |
